# Genome analysis of the unicellular eukaryote *Euplotes vannus* reveals molecular basis for sex determination and tolerance to environmental stresses

**DOI:** 10.1101/357715

**Authors:** Xiao Chen, Yaohan Jiang, Feng Gao, Weibo Zheng, Timothy J. Krock, Naomi A. Stover, Chao Lu, Laura A. Katz, Weibo Song

## Abstract

As a model organism in studies of cell and environmental biology, the free-living and cosmopolitan ciliated protist *Euplotes vannus* has more than ten mating types (sexes) and shows strong resistance to environmental stresses. However, the molecular basis of its sex determination mechanism and how the cell responds to stress remain largely unknown. Here we report a combined analysis of *de novo* assembled high-quality macronucleus (MAC; i.e. somatic) genome and partial micronucleus (MIC; i.e. germline) genome of *Euplotes vannus*. Furthermore, MAC genomic and transcriptomic data from several mating types of *E. vannus* were investigated and gene expression levels were profiled under different environmental stresses, including nutrient scarcity, extreme temperature, salinity and the presence of free ammonia. We found that *E. vannus*, which possesses gene-sized nanochromosomes in its MAC, shares a similar pattern on frameshifting and stop codon usage as *Euplotes octocarinatus* and may be undergoing incipient sympatric speciation with *Euplotes crassus*. Somatic pheromone loci of *E. vannus* are generated from programmed DNA rearrangements of multiple germline macronuclear destined sequences (MDS) and the mating types of *E. vannus* are distinguished by the different combinations of pheromone loci instead of possessing mating type-specific genes. Lastly, we linked the resilience to environmental temperature change to the evolved loss of temperature stress-sensitive regulatory regions of HSP70 gene in *E. vannus*. Together, the genome resources generated in this study, which are available online at *Euplotes vannus* DB (http://evan.ciliate.org), provide new evidence for sex determination mechanism in eukaryotes and common pheromone-mediated cell-cell signaling and cross-mating.

## 1. INTRODUCTION

Single-celled, ciliated protists are abundant in diverse habitats across the globe, where they are among the most important components of food webs in aquatic ecosystems (Lynn 2009). Ciliate diversity, physiology and abundance has been linked to studies of environmental change (Gong et al., 2005; Jiang & Morin 2004; Xu et al., 2014), pollution monitoring (Gutiérrez et al., 2003; Jiang et al., 2011; Stoeck et al., 2018; Yeomans et al., 1997), biogeography (Borror 1980; Foissner 2006; Foissner et al., 2008; Liu et al., 2017; Petz et al., 2007), adaptive evolution (Clark & Peck 2009; La Terza et al., 2004; La Terza et al., 2001; La Terza et al., 2007; Luo et al., 2017; Wang et al., 2017a; Yan et al., 2017) and epigenetics (Wang et al., 2017b; Wang et al., 2017c; Xiong et al., 2016). *Euplotes* is a genus of free-living marine ciliates that play important roles as both predators of microalgae and preys of multicellular eukaryotes like flatworms (Dhanker et al., 2013). For decades, euplotids, including *Euplotes vannus*, have been widely used as model organisms in studies of predator/prey relationships (Kuhlmann 1994; Kuhlmann & Heckmann 1994; Kusch 1993a, b, 1995; Kusch & Kuhlmann 1994; Wiackowski & Szkarlat 1996), cell signaling (Hadjivasiliou et al., 2015; Jerka-Dziadosz et al., 1987), toxicology of marine pollutants (Persoone & Uyttersprot 1975; Trielli et al., 2007) and experimental ecology (Day et al., 2017; Walton et al., 1995; Xu et al., 2004). For example, a previous molecular study revealed that *Euplotes* species have a large number of genes requiring +1 frameshifts (i.e. addition of base pairs post-transcriptional) for expression at the post-transcriptional level, which is much higher than viruses, prokaryotes and other eukaryotes (Wang et al., 2016).

Ciliates, including the models *Paramecium* and *Tetrahymena*, have been shown to present a wide variety of mating type numbers and modes of inheritance (Orias et al., 2017). *Paramecium* has two mating types and its sex determination (SD) is controlled by scnRNA-dependent excision of the SD gene promoter (Singh et al., 2014). *Tetrahymena* has seven mating types, and different mating types specificities are encoded in the single pair of mating type genes in the MAC from all the six pairs in the MIC (Cervantes et al., 2013). More than ten mating types have been identified in *Euplotes* (Akada 1985; Dini & Luporini 1985; Heckmann & Kuhlmann 1986; Kimball 1942; Nanney et al., 1955; Siegel & Larison 1960), yet the molecular basis and SD mechanism are unknown. Previous studies identified two subfamilies of mating-type specific pheromones in euplotids, a shared” pheromone (designated Ec-alpha and named as Type-I pheromone in this work) and a mating type-specific compositional subfamily (designated Ec-1, Ec-2 and Ec-3, and named as Type-II pheromone) (Alimenti et al., 2011; Vallesi et al., 2014). They are considered as the key mediator during the cell-cell signaling that regulates cross-mating processes by controlling self/nonself recognition (Di Giuseppe et al., 2011; Vallesi et al., 2016). Previous studies reported that the euplotid model implicates relationships of hierarchical (or serial) dominance among the pheromone alleles (Dini & Nyberg 1993; Heckmann 1964; Nobili et al., 1978). Yet the results from a recent work indicated that these pheromone genes were expressed without relationships of hierarchical dominance (i.e. heterozygous genotypes behaving like homozygous cells) in the *Euplotes* MAC genome (Vallesi et al., 2014).

Euplotids also feature a strong tolerance to environmental stresses. *Euplotes* spp. were reported to have a conserved molecular defense mechanism to heavy metal contamination for homeostasis by modulating mRNA expression (Kim et al., 2018). In contrast, *E. crassus* and *E. focardii* had a barely detecTable inducible HSP70 response to salinity and temperature stresses (Kim et al., 2017; La Terza et al., 2001). These findings add weight to the argument that the lack of the classical heat shock response might be an adaptation strategy of euplotids to extreme environmental stresses. Additional studies reported *E. vannus*, as a microzooplanktonic grazer, had considerably strong tolerance to ammonia, which may enable it to survive in intensive aquaculture ecosystems with high levels of ammonium, potentially causing great damage in microalgal industry (Day et al., 2017; Xu et al., 2004). Thus, it is important to elucidate the molecular mechanism of *Euplotes* cell response to external stresses under the background of global warming.

In this work, we analyze genomic and transcriptomic data of different mating types of *E. vannus* to study its genomic features, which lead us to reveal the molecular basis of sex determination in euplotids. Furthermore, the gene expression profiling of *E. vannus* cells under different environmental stresses allows us to evaluate how this species tolerates the varying harsh conditions that it encounters.

## 2. METHODS

### 2.1. Cell culture

Six mating types of *Euplotes vannus (*EVJ, EVK, EVL, EVM, EVP and EVX) were collected from seawater along the coast of Yellow Sea at Qingdao (36°06′ N, 120°32′ E), China. Cells of each mating type were cultured separately in filtered marine water at 20°C for 10 days, with a monoclonal population of *Escherichia coli* as the food source, until reaching 10^6^ cells.

### 2.2. Experimental treatment simulating environmental stresses

To simulate the stress from nutrient scarcity, 10^6^ cells of each mating type (EVJ, EVK, EVL, EVM, EVP and EVX) of *E. vannus* were starved for 48 hours before harvest. For stresses from low and high temperature, 10^6^ cells of *E. vannus* mating type EVJ were cultured under the temperature of 4 °C and 35 °C, respectively, for 6 hours before harvest. For stress from low and high salinity, 10^6^ cells of mating type EVJ were cultured under the salinity of 10 psu and 60 psu, respectively, for 6 hours before harvest. For stress from the presence of free ammonia, 10^6^ cells of mating type EVJ were cultured in filtered marine water with 100 mg/L NH_4_Cl (pH 8.3, 20 °C and 35 psu), as described in the previous study (Xu et al., 2004). Cells in two negative control groups were cultured under pH 7.8 and pH 8.2, respectively, in filtered marine water under 20 °C and 35 psu. Each group had two biological replicates.

### 2.3. High-throughput sequencing and data processing

For regular genomic and transcriptomic sequencing to acquire macronucleus (MAC) genome information, cells were harvested by centrifugation at 300 g for 3 min. The genomic DNA was extracted using the DNeasy kit (QIAGEN, #69504, Germany). The total RNA was extracted using the RNeasy kit (QIAGEN, #74104, Germany) and digested with DNase. The rRNA fraction was depleted using GeneRead rRNA Depletion Kit (QIAGEN, #180211, Germany).

For single-cell whole-genome amplification to acquire micronucleus (MIC) genome information, a single vegetative cell of the mating type EVJ of *E. vannus* was picked and washed in PBS buffer (without Mg^2+^ or Ca^2+^) and its MIC genomic DNA was enriched and amplified by using REPLI-g Single Cell Kit (QIAGEN, #150343, Germany), which was based on the whole-genome amplification (WGA) technology and tended to amplify longer DNA fragments.

Illumina libraries were prepared from MAC genomic DNA, mRNA and amplified single-cell MIC genomic DNA of *E. vannus* according to manufacturer’s instructions and paired-end sequencing (150 bp read length) was performed using an Illumina HiSeq4000 sequencer. The sequencing adapter was trimmed and low-quality reads (reads containing more than 10% Ns or 50% bases with Q value <= 5) were filtered out.

### 2.4. Genome assembly and annotation

Genomes of four mating types (EVJ, EVK, EVL and EVM) were assembled using SPAdes v3.7.1 (-k 21,33,55,77), respectively (Bankevich et al., 2012; Nurk et al., 2013). Mitochondrial genomic peptides of ciliates and genome sequences of bacteria were downloaded from GenBank as BLAST databases to remove contamination caused by mitochondria or bacteria (BLAST E-value cutoff = 1e-5). CD-HIT v4.6.1 (CD-HIT-EST, -c 0.98 -n 10 -r 1) was employed to eliminate the redundancy of contigs (with sequence identity threshold = 98%) (Fu et al., 2012). Poorly supported contigs (coverage < 5 and length < 300 bp) were discarded by a custom Perl script.

A final genome assembly of *E. vannus* was merged from the genome assemblies of four mating types (EVJ, EVK, EVL and EVM) by CAP3 v12/21/07 (Huang & Madan 1999). Completeness of genome assembly was evaluated based on expectations of gene content by BUSCO v3 (dataset "Alveolata”) and the percentage of both genomic and transcriptomic reads mapping to the final assembly by HISAT2 v2.0.4 (Kim et al., 2015; Simao et al., 2015). Reads mapping results were visualized on GBrowse v2.0 (Stein 2013). Genome assemblies of *E. crassus* (accession numbers: GCA_001880385.1) and *E. octocarinatus* (accession numbers: PRJNA294366) and their annotation information were acquired from NCBI database and the previous studies (Lobanov et al., 2017; Wang et al., 2018; Wang et al., 2016).

Telomeres were detected by using a custom Perl script which recognized the telomere repeat 8-mer 5’-(C4A4)n-3’ at the ends of contigs, as described in a previous study (Swart et al., 2013). The repeats in the merged genome assembly were annotated by combining *de novo* prediction and homology searches using RepeatMasker (-engine wublast -species ’*Euplotes vannus*’ -s -no_is) (Tarailo-Graovac & Chen 2009). *De novo* genome-wide gene predictions were performed using AUGUSTUS v3.2.2 (--species = euplotes, modified from the model "tetrahymena”, rearranging TAA/TAG as stop codon, TGA as Cys) (Stanke et al., 2006). ncRNA genes were detected by tRNAscan-SE v1.3.1 and Rfam v11.0 (Burge et al., 2013; Lowe & Eddy 1997).

### 2.5. Gene modeling and functional annotation

After mapping RNA-seq data of each mating type of *E. vannus* back to the merged reference genome assembly, the transcriptome of six mating types as well as a merged transcriptome were acquired by using StringTie v1.3.3b (Pertea et al., 2015). Annotation of predicted protein products were matched to domains in Pfam-A database by InterProScan v5.23 and ciliate gene database from NCBI GenBank by BLAST+ v2.3.0 (E-value cutoff = 1e-5) (Camacho et al., 2009; Jones et al., 2014).

### 2.6. Comparative genomic analysis

BLAST+ v2.3.0 was employed to search ciliate gene database from NCBI GenBank to identify corresponding homologous sequences in euplotids (E-value cutoff = 1e-1 and match length cutoff = 100 nt) (Camacho et al., 2009). Joint chromosomes were detected by using a custom Perl script which recognized the chromosomes containing multiple genes (cutoff of distance between two genes = 100 nt). Frameshifting events were detected by using a custom Perl script which recognized the frame change between two BLASTX hits (E-value cutoff = 1e-5 and inner distance <= 10 nt), modified from the protocol in a previous study (Wang et al., 2016), with the addition of a strict criterion (the distance between two adjacent hits with different frames <= 10 bp) to make sure no intron was involved. 30 bp sequences from the upstream and downstream of each type of frameshifting site (+1, +2 or -1) were extracted to identify the motif. Local motifs of nearby frameshifting sites were illustrated by WebLogo 3 (Crooks et al., 2004). The frequency of stop codon usage was estimated by a custom Perl script which recognized the stop codon TAA or TAG in transcripts of euplotids.

### 2.7. Differential gene expression analysis

Transcript abundances were estimated and differential gene expression was analyzed by using featureCounts (Liao et al., 2013) and R packages "Ballgown” and "DESeq2” (p.adjust < 0.01) (Frazee et al., 2015; Love et al., 2014). Starvation induced genes were defined as the average value of RPKM of gene expression from starved samples > 1 and the average value of RPKM of gene expression from vegetative samples < 0.1. Mating type-specific transcripts were defined as the average value of RPKM of gene expression from starved samples > 5 and the average value of RPKM of gene expression from vegetative samples < 0.1. Weighted gene co-expression eigengene network analysis was performed by WGCNA (Langfelder & Horvath 2008). Gene Ontology (GO) term enrichment analysis was performed by using BiNGO v3.0.3 (p.adjust < 0.05), which was integrated in Cytoscape v3.4.0, and the plot was generated by the R package, ggplot2 (Kohl et al., 2011; Maere et al., 2005; Wickham 2016).

### 2.8. Homolog detection of pheromone genes and environmental stress-related genes

Homologous pheromone gene sequences in *E. vannus* were acquired by using BLAST+ v2.3.0 (E-value cutoff = 1e-5), according to the pheromone sequences of *E. crassus* (Alimenti et al., 2011; Vallesi et al., 2014). Genomic DNA samples were harvested from vegetative cells of six mating types of *E. vannus*. Type-II pheromone loci in MAC were amplified using Q5 High-Fidelity 2X Master Mix (NEB, #M0492S, US) with 10 cells of each mating type and genotyping primers (PCR annealing temperature was 64.5 °C, sequences of genotyping primers see Table S9).

Homologous HSP70 gene sequences in *E. vannus* was acquired by using BLAST+ v2.3.0 (E-value cutoff = 1e-5), according to the Hsp70 protein sequences of *E. focardii* and *E. nobilii* from the previous studies (GenBank accession number: AAP51165 and ABI23727, respectively) (La Terza et al., 2001; La Terza et al., 2007). The complete sequences of the *E. focardii* and *E. nobilii* HSP70 genes are available at NCBI with the accession numbers AY295877 and DQ866998, according to the previous studies (La Terza et al., 2004; La Terza et al., 2007). The essential amino acid positions of Hsp70 were reported in previous studies (Morshauser et al., 1999; Sriram et al., 1997). The consensus amino acids sequence of Hsp70 was according to the previous reports (La Terza et al., 2004; La Terza et al., 2007).

### 2.9. Phylogenetic analysis

The DNA and amino acid sequences of *Euplotes* pheromones homologous genes were acquired from NCBI, according to the previous work (Vallesi et al., 2014), and aligned by MUSCLE v3.8.31 and ClustalW v2.1, respectively (Chenna et al., 2003; Edgar 2004). Maximum Likelihood tree based on amino acid sequences was reconstructed by MEGA v7.0.20, using the LG model of amino acid substitution, 500 bootstrap replicates (Kumar et al., 2016; Le & Gascuel 2008).

For phylogenomic analysis by supertree approach, predicted protein sequences of *Euplotes vannus* by us and 31 other ciliates from previous works or transcriptome sequencing by the Marine Microbial Eukaryote Transcriptome Sequencing Project (data available on iMicrobe: http://imicrobe.us/, accession number and gene ID see Table S10) (Aeschlimann et al., 2014; Gentekaki et al., 2017; Keeling et al., 2014; Slabodnick et al., 2017; Wang et al., 2018) were used to generate the concatenated dataset. Maximum Likelihood tree based on the concatenated dataset covering 157 genes was reconstructed by using GPSit v1.0 (relaxed masking, E-value cutoff = 1e-10, sequence identity cutoff = 50%) (Chen et al., 2018) and RAXML-HPC2 v8.2.9 (on CIPRES Science Gateway, LG model of amino acid substitution + Γ distribution + F, four rate categories, 500 bootstrap replicates) (Stamatakis 2014). Trees were visualized by MEGA version 7.0.20 (Kumar et al., 2016).

## 3. RESULTS

### 3.1. General description of genome sequencing and assembly of *Euplotes vannus*

In the current work, we acquired the MAC genomic data of four different mating types (EVJ, EVK, EVL and EVM), which were experimentally confirmed (Table S1). The MAC genome assemblies of these mating type had an average size of 164.2 Mb with a mean coverage of 61X (Table 1 and Figure S1 and Table S2).

**Table 1.** MAC genome assembly and transcriptome-improved gene annotation of *Euplotes vannus* in comparison with that of other euplotids.

|  | *E. vannus* | *E. crassus* | *E. octocarinatus* |
| --- | --- | --- | --- |
| Genome size (Mb) | 85.1 | 58.6 | 88.9 |
| %GC | 37.0 | 38.7 | 28.2 |
| Contig # | 38245 | 56587 | 41980 |
| Contig N50 (bp) | 2685 | 1581 | 2947 |
| 2-telomere contig # | 25519 | 13783 | 29532 |
| 1-telomere contig # | 7835 | 20646 | 4842 |
| 0-telomere contig # | 4890 | 22158 | 7606 |
| 2-telomere contig percentage (%) | 66.7 | 24.4 | 70.3 |
| Genome size (with telomere) (Mb) | 75.9 | 44.9 | 83.1 |
| Scaffold (with telomere) # | 33354 | 34429 | 34374 |
| %Scaffold (with telomere) | 87.2 | 60.8 | 81.9 |
| Scaffold N50 (bp) | 2714 | 1830 | 2999 |
| Gene # | 32755 | - | 29076 |
| Exon # | 175735 | - | 96843 |
| Transcript # | 43040 | - | 29076 |

After the contigs identified as noise (coverage < 5X) or contamination of bacteria and mitochondria were removed, the final genome assembly of *E. vannus* was generated by merging the genome assemblies of these four mating types (assembled genome size is 85.1 Mb and N50 = 2,685 bp, Table 1). We compared these data with those of two other euplotids, *E. crassus* and *E. octocarinatus*. Although *E. vannus* and *E. crassus* shared a similar %GC (36.95% and 38.65%, respectively), the genome size, number of 2-telomere contigs and N50 value of *E. vannus* were more comparable with those of *E. octocarinatus* (Table 1).

The contig N50 values of these three *Euplotes* species were all smaller than 3 kb, because chromosomes of *Euplotes* species are "nanochromosomes ", similar to that of *Oxytricha trifallax* (Wang et al., 2016). The 2-telomere contig percentage of the merged genome of *E. vannus* is 66.7%, consistent with *E. octocarinatus* in which 70.3% contigs contained telomeres on both ends. Among the four mating types with genomic sequencing data, the genome assembly of EVJ was of highest quality with a 2-telomere contig percentage of 81.6% and N50 of 2,954 bp (Table S2).

To evaluate the completeness of the genome assembly of *E. vannus*, gene content from single-copy orthologs of protists was identified by BUSCO, and the result indicated that the current assembly had a comparable percentage of complete ortholog sequences with other species (Figure 1 and Figure S2). Furthermore, the majority of genomic DNA sequencing reads of four mating types (EVJ, EVK, EVL and EVM) and RNA-seq reads of six mating types (EVJ, EVK, EVL, EVM, EVP and EVX) in both starvation and vegetative stages can successfully be mapped back to the merged reference genome assembly with a mean mapping ratio of 80.1% (Table S3). Furthermore, 109 tRNAs which consist of 48 codon types for 20 amino acids, were detected in the final genome assembly (Table S4). These results indicated that our genome assembly of *E. vannus* was largely complete. The information of repeat regions and functional annotation of genes are summarized in Figure S3, Table S5 and Table S6.

**FIGURE 1.**
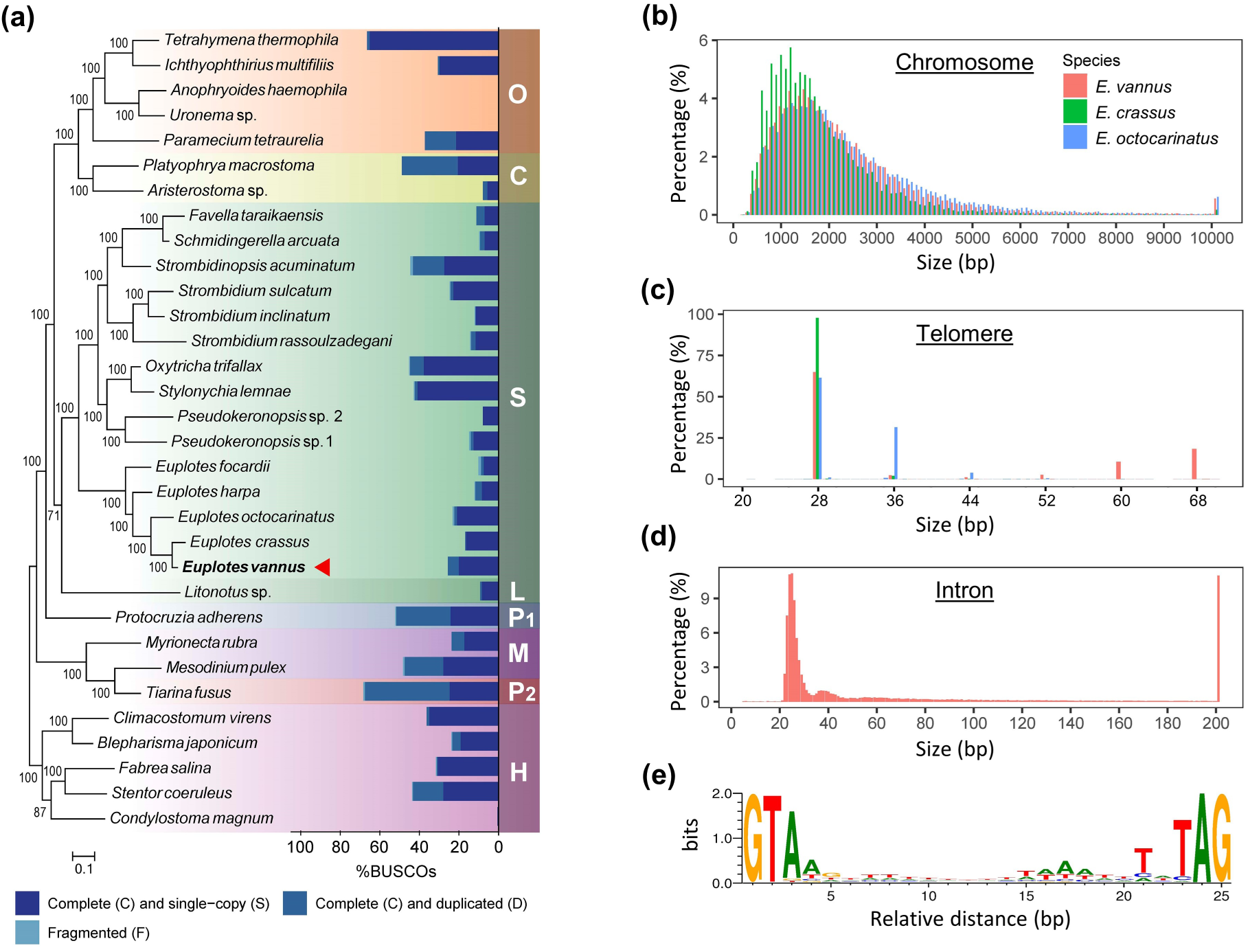
Genome assembly of *E. vannus*. (a) Maximum likelihood phylogenetic tree by supermatrix approach and assembly completeness evaluation of ciliate genomes/transcriptomes by BUSCO. Dark blue blocks represent the percentage of complete and single-copy genes among protists, and the steel-blue blocks represent that of complete and duplicated genes in each species. Genomic data of *Euplotes vannus* sequenced in the current work is marked by the red triangle. S: class Spirotrichea. L: class Litostomatea. O: class Oligohymenophorea. C: class Colpodea. P1: class Protocruzia. M: class Mesodiniea. P2: class Prostomatea. H: class Heterotrichea. The scale bar corresponds to 10 substitutions per 100 nucleotide positions. (b) Size distribution of 2-telomere scaffolds of *E. vannus, E. crassus* and *E. octocarinatus*. (c) Size distribution of telomeres of *E. vannus, E. crassus* and *E. octocarinatus*. (d) Size distribution of introns in *E. vannus* mating type EVJ. (e) Sequence motif of 8792 tiny introns with the size of 25 nt. Weblogo was generated and normalized to neutral base frequencies in intergenic regions.

The size distribution of complete chromosomes of *E. vannus* (i.e. those bearing telomeric repeats "C4A4" and "T4G4" on both ends) is quite close to that of another two euplotids, *E. octocarinatus* and *E. crassus*, with the peak values around 1.5 kb (Figure 1B). A similar result was found in the size distribution of telomeres, in which most telomeres of all three euplotids had a length of 28 bp, with an increment of 8 bp (Figure 1C). Most identified *E. vannus* introns were around 25 bp in length, with a canonical sequence motif 5’-GTR (N)nYAG-3’ at either end (Figure 1D, E and Figure S4).

### 3.2. Evolution and synteny/comparative genomic analyses among euplotids

Most chromosomes (37501/38245, 98.1%) in *E. vannus* are nanochromosomes, or so-called gene-sized chromosomes, bearing a single gene on each. A similar result was observed in *E. octocarinatus* (40396/41980, 96.2%). There was a small proportion of chromosomes that contains more than one gene (Figure 2A). These joint nanochromosomes were then divided into two groups by the consistency and inconsistency of the transcription directions of the genes on them (cis and trans, respectively). The trans-joint nanochromosomes had close numbers, considering the total chromosome numbers were similar in these two euplotids (Table 1). However, *E. octocarinatus* possessed 2-fold more cis-joint nanochromosomes than *E. vannus*.

**FIGURE 2.**
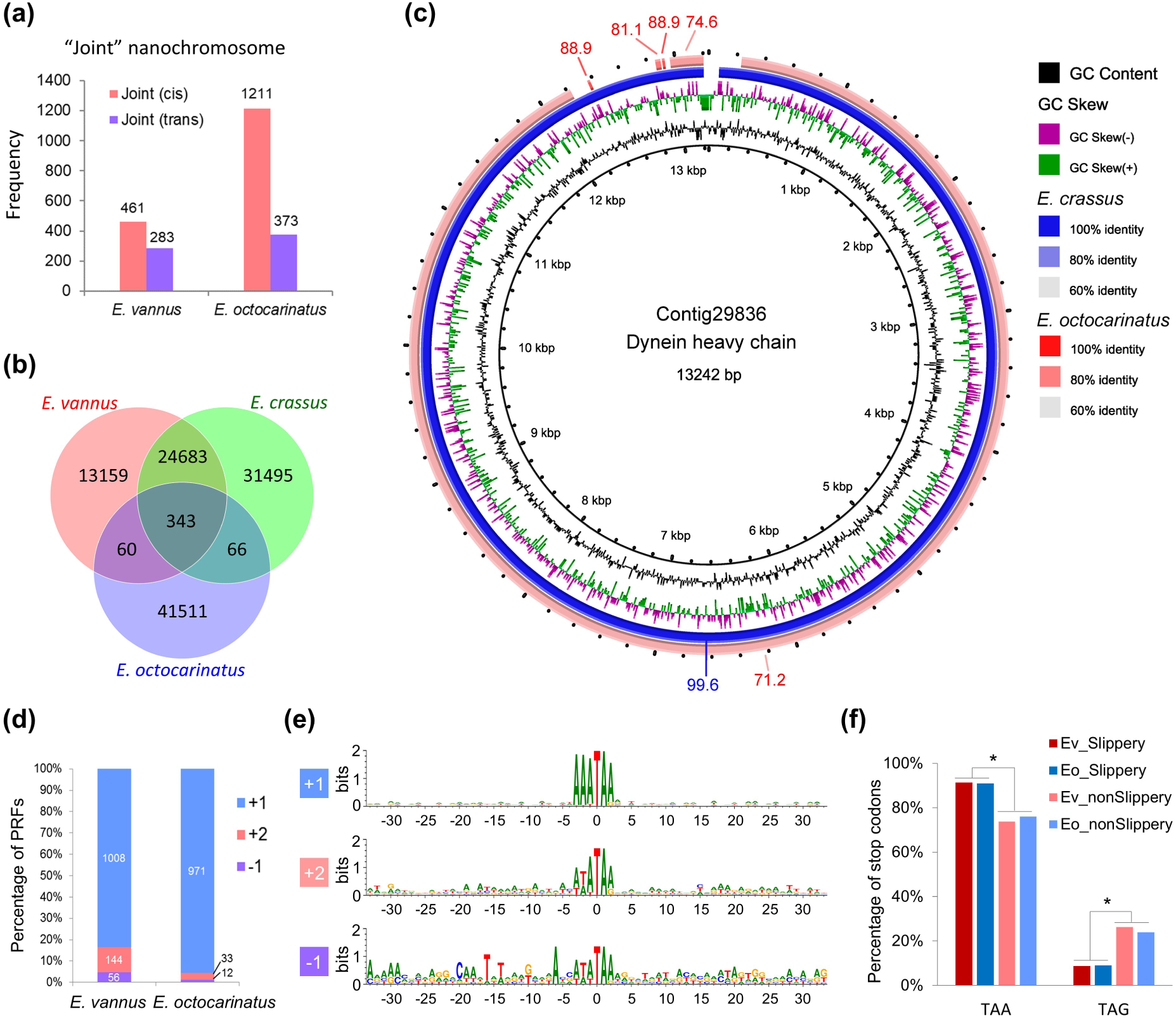
Evolution and synteny/comparative genomic analyses. (a) Joint nanochromosome detection in *E. vannus* and *E. octocarinatus*. Red bars denote the frequency of joint nanochromosomes containing genes in a same transcription direction (cis) and purple bars denote that of joint nanochromosomes containing genes in opposite transcription directions (trans). (b) Closely related contigs among three euplotids. (c) Homologous comparison of the contigs containing dynein heavy chain coding gene among *E. vannus* (as reference), *E. crassus* (blue) and *E. octocarinatus* (red). (d) Frameshifting detection & comparison with *E. octocarinatus*. (e) Conserved sequence motif associated with frameshift sites. Sizes of letters denote information content, or sequence conservation, at each position. The analysis is based on the alignment of 30 bp upstream and downstream the frameshifting motif from 1236 predicted +1 frameshifting events that involves stop codon TAA or TAG. Note the canonical motif 5’-AAA-TAR-3’ (R = A or G) in +1 PRF and noncanonical motif 5’-WWW-TAR-3’ (W = A or T) in +2 and -1 PRF. (f) Stop codon usage in slippery and non-slippery transcripts of *E. vannus* (Ev) and *E. octocarinatus* (Eo). Asterisks denote the significant difference (*p* < 0.01).

*E. vannus* and *E. crassus* shared 25026 closely related contigs (E-value cutoff = 1e-5), which was equivalent to 65.4% and 44.2% of total contigs for each species, respectively (Figure 2B). However, only 469 contigs were shared between these two species and *E. octocarinatus*. Furthermore, *E. vannus* and *E. crassus* shared not only more homologous contigs, but also more sequence identity (Figure 2C). For example, the sequence identity between *E. vannus* and *E. crassus* of the corresponding region on the chromosome that contains the coding gene of dynein heavy chain protein was 99.6%, while the sequence identity between *E. vannus* and *E. octocarinatus* was 71.2%.

Frameshifting events in *E. vannus* and *E. octocarinatus* were detected by identifying the adjacent region between two BLASTX hits that are in different frames, targeting to a same protein sequence (illustrated by Figure S5). The E-value cutoff (1e-5) ensured the accuracy of the prediction process and a small inner distance cutoff (10 nt) was applied to get rid of the interference from introns, because all introns of euplotids were larger than 20 nt as described above (Figure 2D). The result indicated that the high frequency of +1 programmed ribosomal frameshifting (PRF) was a conserved feature in euplotids. However, intriguingly, more +2 and -1 PRF events were found in *E. vannus* (16.6%) than in *E. octocarinatus* (4.4%). With more cases of +2 and -1 PRF events being spotted, a novel motif rather than the canonical motif 5’-AAA-TAR-3’ (R = A or G) was revealed as 5’-WWW-TAR-3’ (W = A or T) (Figure 2E).

Based on the genomic and transcriptomic sequencing data, the stop codon usage was analyzed in *E. vannus* and *E. octocarinatus* and compared between the regular transcripts and the slippery sites of PRF transcripts (Figure 2F). In these two euplotids, UAA was preferentially used in the regular termination signal (73.7% and 76.0%, respectively) and in the slippery signal (91.3% and 91.0%, respectively). Moreover, the frequency of UAA codon usage in slippery signal is significantly higher than that in the regular termination signal (*p* = 0.005024 < 0.01, Analysis of variance), which suggested that UAA may be favorable for frameshifting in both *E. vannus* and *E. octocarinatus*.

### 3.3. Profiling and development of *Euplotes* pheromone genes

Pheromone alleles were successfully identified in MAC genomes of four *E. vannus* mating types (Table S7). Other than the pheromones Ev-1, Ev-2, Ev-3 and Ev-alpha that were orthologous to Ec-1, Ec-2, Ec-3 and Ec-alpha in *E. crassus*, respectively, a novel Type-II pheromone Ev-4 and a novel Type-I pheromone Ev-beta were found in *E. vannus* (Figure 3A and Figures S6, S7 and S8). Although Ev-4 used "TAG” as stop codon rather than using "TAA” in other three Type-II pheromones (Figure S6), together with Ev-beta, they showed a significant sequence similarity with the other three known pheromones, especially in the pre- and pro-regions of the cytoplasmic precursor (marked by red arrows in Figure 3A) and retained highly conserved cysteine residues in the secreted region (marked by red dots), which was an important feature of pheromone allele sequences.

**FIGURE 3.**
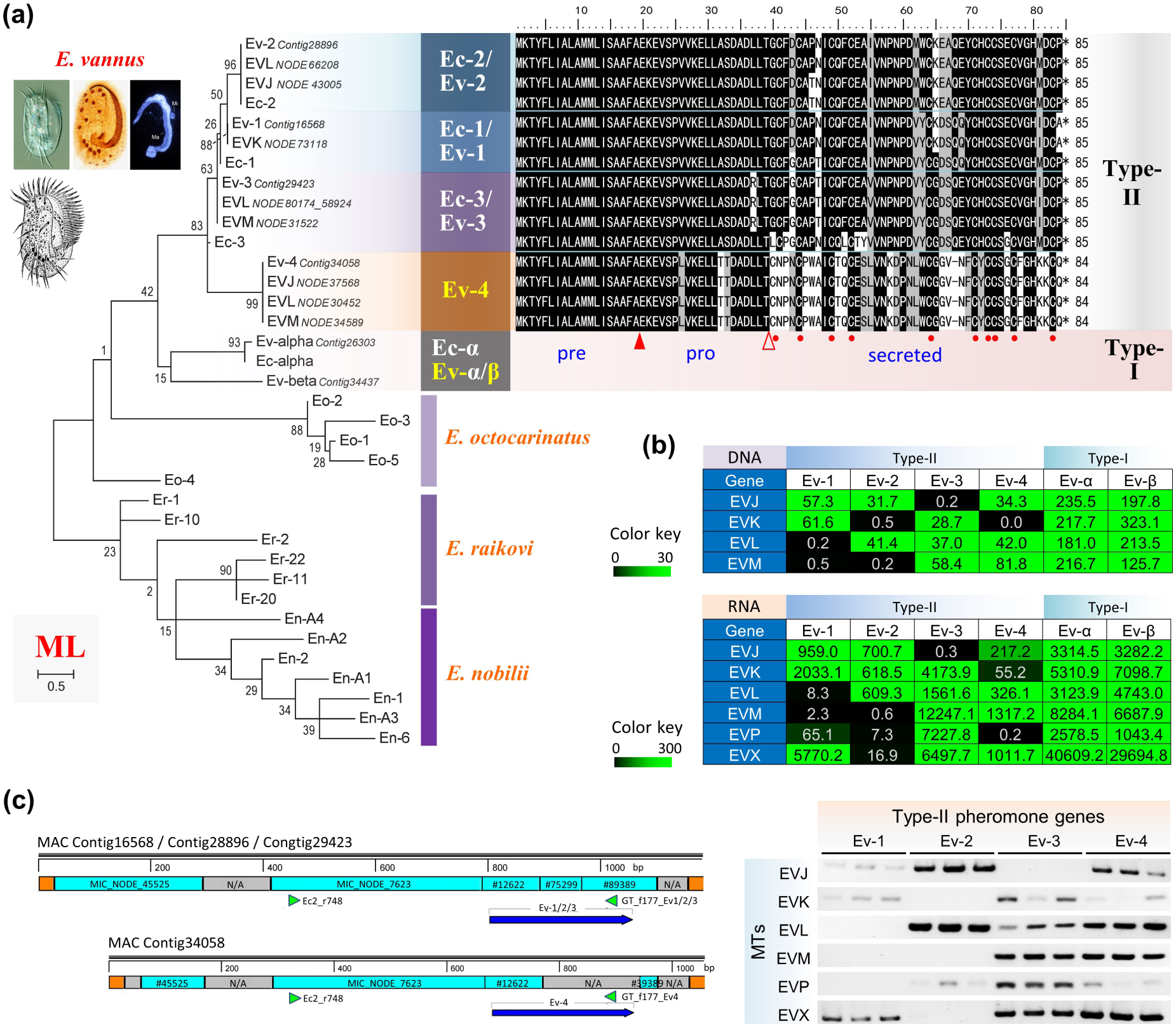
Genomic investigation revealed new pheromone loci Ev-4 and Ev-beta and a mating type-specific combination of these loci in *E. vannus*. (a) Phylogenetic analysis of *Euplotes* pheromones and sequence alignment of Type-II pheromones of *E. vannus* and *E. crassus*. Identical residues are shadowed in black and similar residues are shaded in grey. Asterisks mark the positions of stop codons. Filled and light arrowheads indicate the extension positions of the pre- and pro-regions, respectively. Red dotes denote the 10 conserved cysteine residues in secreted region. Numbers indicate the progressive amino acid positions in the sequences. (b) Chromatin and gene expression profiling based on the RPKM of pheromone loci in each mating type of *E. vannus* by genome and transcriptome sequencing. The Tables are colored by RPKM values. (c) The primer design (left) and PCR amplification results (right) of *E. vannus* Type-II pheromone genes. Boxes in orange, blue and grey denote the telomeres of the MAC contigs containing the pheromone genes, the MDS regions and regions those loci in MIC genome still unknown (N/A), respectively. Arrow heads in green denote the positions of the primers. Arrows in blue denote the coding regions of pheromone genes. PCR amplifications of each pheromone gene in each mating type are conducted with three biological replicates.

The phylogenetic analysis of *Euplotes* pheromones, including the homologs in each mating type of *E. vannus* and the corresponding consensus sequences, was performed (Figure 3A). Although these MAC loci were generated by alternative processing of MIC regions in each species, the result indicated that the pheromones of *E. vannus* clustered together with those of *E. crassus*, distinct from those of other Euplotids, *E. octocarinatus, E. nobilii* and *E. raikovi*.

Chromatin profiling of pheromone genes indicated that the combination of Type-II pheromone genes, Ev-1, Ev-2, Ev-3 and Ev-4, exhibited a mating type-specific feature on genic level (Figure 3B and Figure S9). Different mating types retained 1-3 Type-II pheromone genes in the MAC genome and no duplicate events existed. Gene expression profiling of pheromone genes consistently showed that both two Type-I pheromones Ev-alpha and Ev-beta were highly expressed in all six mating types in *E. vannus* and confirmed the mating type-specific chromatin profiling of Type-II pheromone genes in different mating types at the transcriptional level. As another independent verification, PCR amplification of pheromone loci was carried out to verify the presence of the pheromone-related contigs in the MAC genome of each mating type (Figure 3C). The results of PCR amplification of pheromone loci indicated that Type-II pheromone gene Ev-1 was absent from the mating types EVL, EVM and EVP, Ev-2 was absent from EVK, EVM, EVP and EVX, Ev-3 was absent from EVJ and Ev-4 was absent from EVK and EVP, which were mostly consistent with the results from chromatin and gene expression profiling (Figure 3B). In brief, each of the six mating types we identified contained a unique combination of four Type-II pheromone genes.

To further study the development process of the pheromone genes during the programmed DNA rearrangement from germline MIC to somatic MAC in *E. vannus*, MIC genomic DNA was acquired by single-cell sequencing and its draft genome was assembled (Table 2). Then the germline genome (MIC) origins of Ev-1 (MAC Contig16568), Ev-2 (Contig28896), Ev-3 (Contig29423) and Ev-4 (Contig34058) were mapped to MIC contigs (Figure 3C and Table S8, E-value cutoff = 1e-5). The coding region of pheromone genes Ev-1, Ev-2 and Ev-3 consisted of three MDS regions from the MIC genome while Ev-4 consisted of at least two MDS regions. However, the germline source of a part of the pro-region and the secreted region of Ev-4 was not found.

**Table 2.**
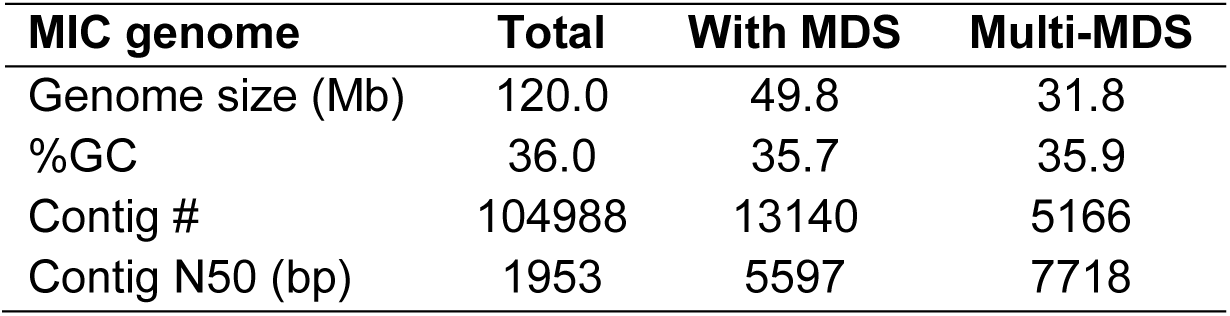
MIC genome assembly information of *Euplotes vannus* and recognition of MDS-containing contigs and those that contain multiple MDSs.

### 3.4. Molecular basis of strong tolerance to extreme environmental stresses

Starvation stress (i.e. nutrient scarcity) had limited impact on the expression pattern of different mating types by global transcription profile (Figure 4A and Figure S10). However, differential gene expression analysis revealed some mating type-specific transcripts under nutrient scarcity (Figure 4B). Transcripts induced by starvation tended to be associated with mating type-specific genes (42.8%; see Figure S11). Gene functional annotation reveals that these starvation-induced and mating type-specific transcripts are related to protein transport and phosphorylation process in cells (Figure 4C and Figure S12). The expression of these genes may facilitate the cell response to pheromone-mediated cell-cell signaling and cross-mating behavior under the stress from nutrient scarcity.

**FIGURE 4.**
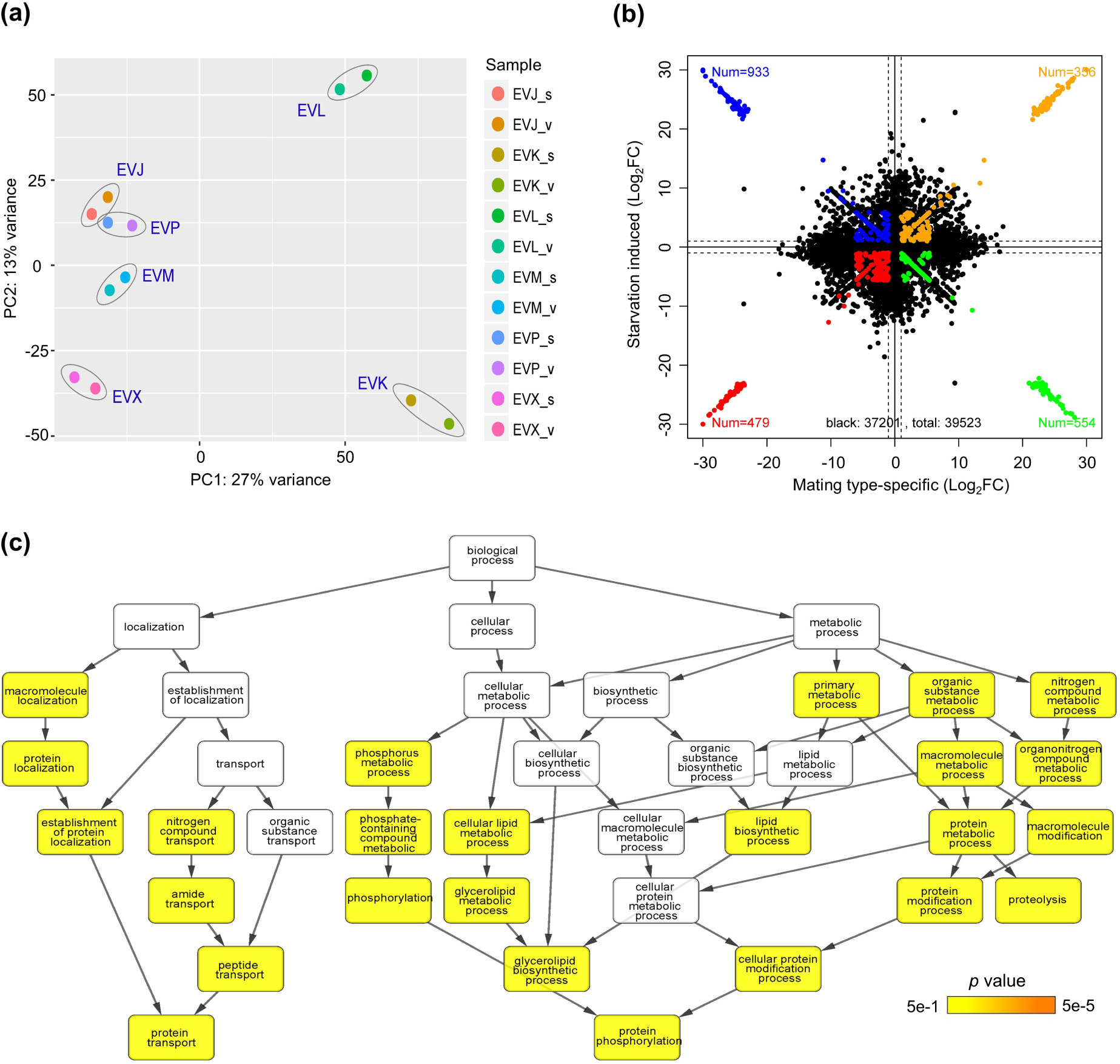
Gene function analysis of mating type-specific transcripts of *E. vannus* indicates they are starvation-induced and related to protein transport and phosphorylation process in cells. (a) PCA analysis on transcript enrichment of different mating types. (b) Cross plot of differential expression of mating type-specific and starvation-regulated transcripts. (c) GO enrichment analysis of mating type-specific and starvation induced transcripts.

PCA analysis based on the differential gene expression of *E. vannus* EVJ cells under different extreme environmental stresses was performed (Figure S13A). The result revealed changes in the gene expression profile of cells under high temperature (35 °C), low temperature (4 °C), high or low salinity (60 and 10 psu, respectively). Also, the presence of free ammonia had substantial impact on transcription patterns (Figure S13). Surprisingly, cells under high salinity and low salinity shared a similar gene expression profile (Figure S13A).

To further dissect the relationships between co-expression of genes associated with the regulation of cellular processes and pathways under extreme environmental changes, a weighted gene co-expression eigengene network was constructed (Figure 5A). The network clustered different eigengenes into six modules based on their co-expression profile (Figure S13B). A strongly co-expressed eigengene module was up-regulated in cells under both high and low salinity stresses (colored in steel blue in Figure 5A and Figure S13B). This module was involved with an extensive activation of many pathways, mainly related to tRNA aminoacylation, tRNA and rRNA processing, nucleosome assembly and pseudouridine synthesis (p.adjust < 0.05). In addition, two small eigengene modules were up-regulated in cells under high salinity stress (purple) and low salinity stress (dark green), respectively, and low salinity stress activated an extra pathway related to the glutamine metabolic process.

**FIGURE 5.**
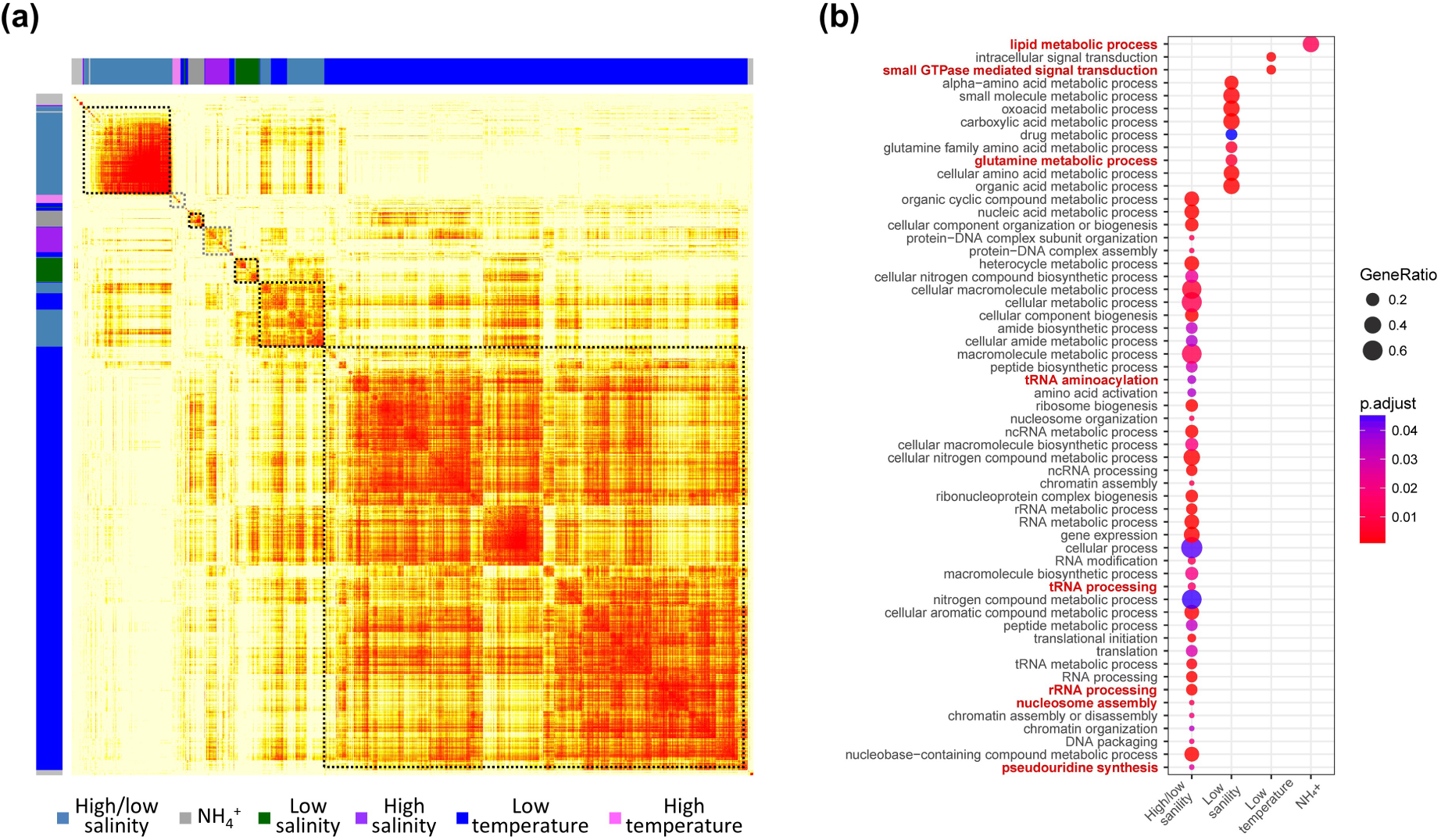
Differential gene expression analysis reveals several large cohorts of co-expressed genes under temperature, salinity and free ammonia stresses. (a) Heatmap of weighted gene co-expression network, in accordance with different stress-response gene groups. (b) GO term enrichment analysis on different stress-response gene groups.

Intriguingly, low temperature stress induced a large module cluster of eigengenes related to small GTPase mediated signal transduction, while very few eigengenes co-expressed under the high temperature (blue and purple, respectively, in Figure 5 and Figure S13B). The homolog of the highly conserved heat-shock protein 70 (Hsp70), which many organisms upregulate under environmental stress, was identified in *E. vannus* and compared with its counterparts in *E. nobilii* and *E. focardii* (Figure S14). The result revealed that only the *E. focardii* Hsp70 sequence had numerous amino acid substitutions within its two major functional domains, i.e. the ATP-binding and substrate-binding domains (Figure 6A). However, the transcription of the HSP70 gene in *E. vannus* did not respond to temperature stresses while being responsive to other stresses like salinity and chemical stresses (Figure 6B).

**FIGURE 6.**
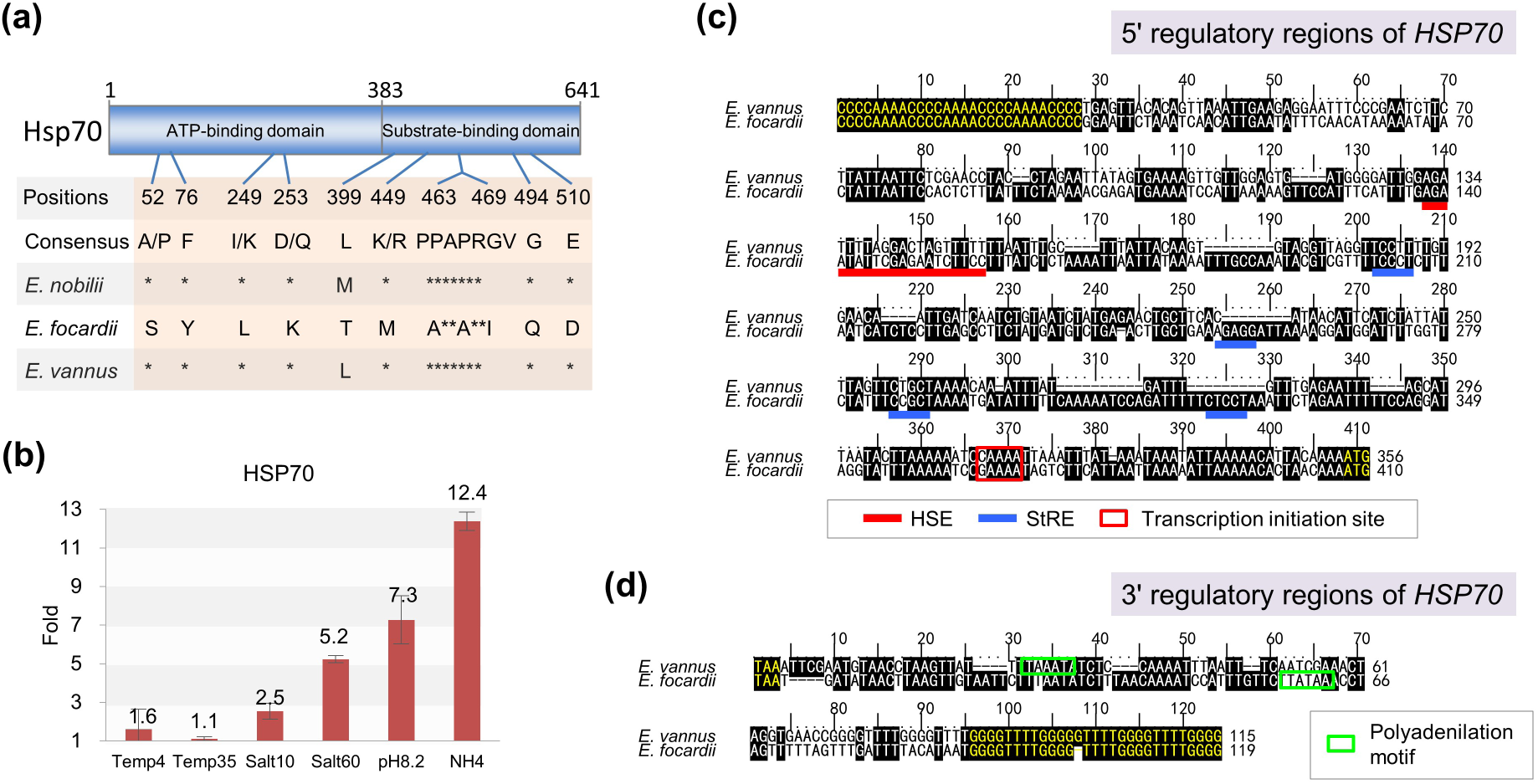
Gene expression and sequence analysis of HSP70 in *E. vannus* shows it has evolved to be insensitive to temperature change. (a) Amino acid substitutions that occur in *E. focardii* at the level of its HSP70 ATP- and substrate-binding domains and are unique with respect to *E. nobilii* and other organisms. Asterisks denote identities. Numbers indicate essential amino acid positions of Hsp70. (b) Up-regulation fold change in cells under different environmental stresses (4 °C, 35 °C, 10 psu, 60 psu, pH 8.2 or with the presence of free ammonia) with respect to the control (25°C, 35 psu and pH 7.8). (c) Nucleotide sequence alignment of the 5’ regulatory regions of the *E. vannus* (gene MSTRG.11315” on contig Contig21532”) and *E. focardii* HSP70 genes. The identities are shaded; the telomeric C4A4 repeats and transcription initiation ATG codons are in yellow; putative sites for the transcription initiation are boxed; sequence motifs bearing agreement with HSE and StRE elements are indicated by red and blue bars, respectively. (d) Nucleotide sequence alignment of the 3’ regulatory regions of the *E. vannus* and *E. focardii* HSP70 genes and neither of them carries ARE elements in the 3’ regulatory region. The identities are shaded; the telomeric G4T4 repeats and stop TAA codons are in yellow; putative polyadenylation motifs are boxed.

To gain a better understanding of the molecular basis of the lack of change in response of the HSP70 gene to temperature stress in *E. vannus*, we analyzed the structure of non-coding regions flanking the gene in *E. vannus* and compared the structure of this gene between *E. vannus* and *E. focardii* (Figure 6C, D). The result indicated that no substantial difference was detected in the 5’ promoter region between the HSP70 genes of these two species, both bearing canonical regulatory *cis*-acting elements that bind transcriptional trans-activating factors, including heat-shock elements (HSE) and stress-response elements (StRE) (Figure 6C). However, the sequence of HSEs in these two species is poorly conserved. Furthermore, neither *E. vannus* nor *E. focardii* retained an mRNA destabilization adenine-rich element (ARE) in their 3’ promoter region (Figure 6D).

In contrast, the gene expression of cells under the presence of free ammonia were very similar to those under high temperature stress, and there was a small cluster of eigengenes that responded related to the lipid metabolic process (Figure 5 and Figure S13B). However, chemical stress activated the expression of HSP70 gene in *E. vannus* significantly (Figure 6B).

## 4. DISCUSSION

### 4.1. Genomes of ciliates are divergent to each other and largely unexplored

In the current work, we present a high-quality macronuclear and a partial micronuclear genome assemblies of a unicellular eukaryote, *Euplotes vannus*, which possesses gene-sized” MAC nanochromosomes. Comparative genome analysis reveals that *E. vannus* shares similar patterns on frameshifting and stop codon usage with *E. octocarinatus* and is undergoing incipient sympatric speciation with *E. crassus* (figs. 2 and 3). Besides euplotids, we also evaluated the assembly completeness of genomes or transcriptomes of other ciliates and found a large divergence among them (Figure 1A and Figure S2). It reflects two facts: 1) ciliates have great genetic distances, even those closely related on phylogeny; 2) ciliate genomic data sequenced so far has not been collected by assembly completeness evaluation tools like BUSCO (the only ciliate covered is *Tetrahymena thermophila*). Nevertheless, among these ciliates, the completeness of transcriptome assembly of three species, *Anophryoides haemophila, Uronema* sp. and *Condylostoma magnum*, were not evaluated. One of the most likely reasons is that they possess small genome volume and thus share few genes with other ciliates (Chen et al., 2018). For instance, *Anophryoides* and *Uronema* are well-known parasitic species that cause disease in fish and lobsters in aquaculture facilities and have very small genome sizes (Acorn et al., 2011; Cheung et al., 1980; Dragesco et al., 1995; Iglesias et al., 2001; Munday et al., 1997). Another reason is that stop codon rearrangement occurs in some ciliates, such as *Condylostoma magnum*, and all standard stop codons are reassigned to amino acids in a context-dependent manner (Heaphy et al., 2016; Swart et al., 2016). These unusual features could dramatically increase the difficulty to precisely evaluate the assembly completeness of ciliate genomes/transcriptomes. Overall, the genomic investigation in ciliates is still waiting to be explored and the evaluation of ciliate genome assemblies calls for further improvement.

### 4.2. Genomic data and pheromone gene assay supports the hypothesis that *E. vannus* and *E. crassus* may be undergoing incipient sympatric speciation

An early study reported that *E. vannus* could interbreed with *E. crassus* under laboratory conditions (Valbonesi et al., 1988). Other studies reported that *E. vannus* and *E. crassus* might be undergoing sympatric speciation (Dobzhansky 1940; Zhao et al., 2018). The result in the current work increased the weight of this argument in three aspects: 1) *E. vannus* and *E. crassus* are closely clustered with each other among euplotid species (Figure 1A); 2) these two species shared both a large number of homologous sequences and at high levels of sequence identity, in contrast to comparisons with *E. octocarinatus* (Figure 2B, C); 3) the homologs of pheromones of *E. vannus* and *E. crassus* are closely related and distinct from those from other *Euplotes* species (Figure 3A, B). Orthologs of each pheromone allele from different mating types of *E. vannus* share identical sequences on both gene and protein levels in most cases except pheromone gene Ev-2 (Figure 3A and Figures S6, S7 and S8). Our findings might describe a pattern that mating type loci evolve rapidly after a recent speciation between *E. crassus* and *E. vannus*.

### 4.3. Molecular basis of the sex determination in *E. vannus*: the combination of Type-II pheromone loci is mating type-specific

Type-I pheromone Ec-alpha has large sequence differences with the Type-II pheromones and has been considered as an adaptor” that interacts with the other pheromones, as it has a strong propensity to oligomerize and retains a hydrophilic domain for putative interaction (Vallesi et al., 2014; Vallesi et al., 2016). This argument is supported by the results of expression profiling for pheromone genes in the current work (Figure 3B).

Studies on the pheromones from other euplotids, including *E. raikovi* (Brown et al., 1993; Liu et al., 2001; Luginbühl et al., 1994; Mronga et al., 1994; Ottiger et al., 1994; Weiss et al., 1995; Zahn et al., 2001), *E. nobilii* (Di Giuseppe et al., 2011; Pedrini et al., 2007; Placzek et al., 2007; Vallesi et al., 2012) and *E. octocarinatus* (Brünen-Nieweier et al., 1998; Kuhlmann et al., 1997; Möllenbeck & Heckmann 1999; Schulze Dieckhoff et al., 1987), revealed that highly enriched and conserved cysteine residues in the secreted region is the most outstanding sequence motif of *Euplotes* pheromones (Vallesi et al., 2016). The novel pheromone Ev-4 identified in this study from *E. vannus* retains 10 cysteines, as same as other Type-II pheromones in *E. vannus* and *E. crassus*, and thus matches this characteristic perfectly (Figure 3A).

As the Ec-1 and the other two Type-II pheromones, Ec-2 and Ec-3, were identified in different mating types of *E. crassus* by pheromone purification and molecular mass determination after chromatographic separation, confirmed by PCR amplification and sequencing, Type-II pheromones have been considered as mating type-specific (Alimenti et al., 2011; Vallesi et al., 2014). However, our study demonstrated that six *E. vannus* mating types retain different combinations of Type-II pheromone loci in their MAC genomes (Figure 3C), and thus they exhibited highly different pheromone gene expression profiling instead of possessing exclusive, mating type-specific genes (Figure 3B). Furthermore, mating types EVL and EVP have the same set of pheromone genes but with different abundance (Figure 3C). Therefore, it suggests that the mating type-specific combination of the Type-II pheromone loci might not be an all-or-none phenomenon, but a manner related to the composition or copy number of Type-II pheromone genes. Although further studies are expected, the observations of mating type-specific combination in the current study support the allelic codominance or non-hierarchical dominance relationship among signaling pheromone genes in euplotids (Vallesi et al., 2014; Vallesi et al., 2016).

Taken together, the current work revealed that euplotids have a novel SD manner by which mating types are determined through mating type-specific combination of four Type-II pheromone genes (Figure 7C). Unlike *Paramecium tetraurelia*, there is no excision event on promoter regions of *E. vannus* pheromone genes (figs. 3C, 7A and Figure S6). On the other hand, the MAC of *E. vannus* does not possess exclusive mating type-specific SD loci as *Tetrahymena thermophila* (Figure 3 and 7B). This mating type-specific feature of Type-II pheromones comes from the programmed DNA rearrangement between germline and somatic genomes. Intriguingly, none of the *E. vannus* mating types we have identified possesses all four Type-II pheromone genes in MAC (Figure 3B and Figure S9). Thus, the results of current study described a third SD type in ciliates (Figure 7C).

**FIGURE 7.**
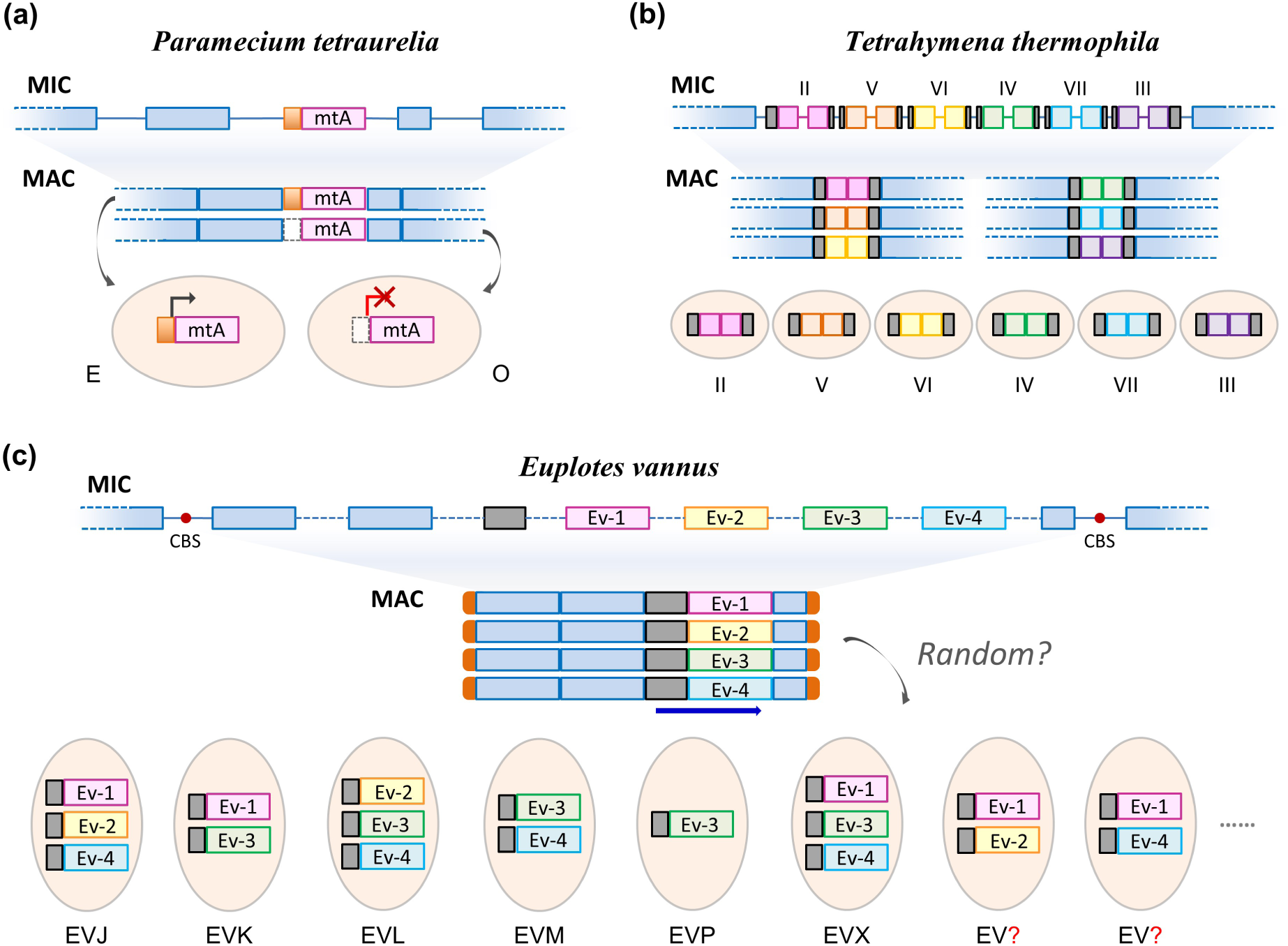
Current models (simplified) for sex determination in *Paramecium tetraurelia, Tetrahymena thermophila* and *Euplotes vannus*. (a) One of the two *P. tetraurelia* mating types, mating type E depends on expression of the mtA gene during sexual reactivity. The other mating type O is determined during macronuclear development by excision of the mtA promoter (box in orange) as an internal eliminated sequence (IES), preventing expression of the gene. Adapted from Extended Figure 10 of Reference (Singh et al., 2014). (b) Mating type gene pairs in *Tetrahymena thermophila* macronuclear are assembled by joining mating type-specific macronucleus destined sequence (MDS) from micronuclear to reproduce six mating types. Segments filled with grey represent conserved transmembrane regions. Adapted from Figure 3 of Reference (Cervantes et al., 2013). (c) *E. vannus* shows a mating type-specific feature on the combination of different pheromone genes in MAC. Red dots denote chromosome breakage site (CBS). Segments filled with solid orange, light colors and grey denote telomeres, MDSs and conserved transmembrane regions, respectively. Dashed lines denote putative IESs.

### 4.4. Molecular basis of the HSP70’s lack of response to temperature stress

A previous study indicated that response of the HSP70 gene expression to temperature change, no matter gradually or abruptly, was divergent between *E. nobilii* and *E. focardii* (La Terza et al., 2001). When transferred from 4 to 20 °C, a strong transcriptional activity of HSP70 genes was induced in *E. nobilii* cells, while no measurable change was found in cells of *E. focardii*. In contrast, HSP70 expression increased with oxidative and chemical stresses such as tributyltin and sodium arsenite (La Terza et al., 2004). Furthermore, together with the results from the previous studies (Dobzhansky 1940), the current work strongly suggests the HSP70 gene of *E. vannus*, which is largely divergent from that of *E. focardii*, does not carry unique amino acid substitutions of potential significance for cold adaptation (Figure 6A).

The HSP70 gene of the cosmopolitan species *E. vannus* was only activated when facing with chemical stresses but not thermal changes (Figure 6B). The study in Antarctic psychrophilic euplotid *E. focardii* also reported similar Hsp70 transcriptional activity which exclusively responded to non-thermal stresses (La Terza et al., 2004). *E. focardii* harbors the *cis*-acting elements like heat-shock elements (HSE) and stress-response elements (StRE) in 5’ promoter region, which are known to be targets of trans-acting transcriptional activators characterized in a variety of organisms in association with their stress-inducible genes (Fernandes 1994; Kobayashi & McENTEE 1993; Ruis & Schüller 1995). It is also argued that the HSE-modulated HSP70 gene transcription is more specific for a response to temperature stress while the StRE-modulated HSP70 gene transcription is more specific for a response to a broader range of non-temperature stresses (La Terza et al., 2007). However, the corresponding HSE element of *E. vannus* is poorly conserved, which may impair its sensitivity to temperature stress (Figure 6C). Furthermore, the absence of an adenine-rich element (ARE) in the HSP70 3’ regulatory region which would exclude a rapid mRNA degradation was also observed in *E. vannus* (Figure 6D), as reported in the previous study of *E. focardii* (La Terza et al., 2007). Combining these data with observations in the current work, we argue that structural divergence of transcriptional *trans*-activating factors underlies that lack of change of HSP70 gene expression in response to temperature stress in *E. vannus* and *E. focardii*.

## 5. CONCLUSION

In summary, the results of the current study indicate that *Euplotes vannus* has a set of orthologous pheromones to the reported ones in *E. crassus*, as well as a novel type of pheromone named as Ev-4 which also shares close homology with *E. crassus*, and thus explains the hybridization between these two species on the molecular level. Chromatin and expression profiling of pheromone genes indicated that the combination of these genes is mating type-specific on the genic level. Our work provides new evidence for sex determination mechanism and supports the allelic codominance or non-hierarchical dominance relationship among pheromone loci during *Euplotes* pheromone-mediated cell-cell signaling and cross-mating. Furthermore, the homologous search between MAC and MIC genomes reveals that pheromone genes in *E. vannus* develop by programmed DNA rearrangement. According to the analyses of transcriptomes under different environmental stresses, although the HSP70 gene of *E. vannus* does not carry unique amino acid substitutions of potential significance for cold adaptation, it has evolved to be insensitive to temperature change by losing the temperature stress-sensitive HSE element and mRNA destabilization ARE elements in the regulatory region of HSP70.

## ACKNOWLEDGEMENTS

The authors would like to thank the following people for assistance with this study: Tengteng Zhang and Ruitao Gong (Ocean University of China, China), for the assistance of experimental verification; Dr. Henglong Xu (Ocean University of China, China), for the advice on the experiment design of environmental stresses; Dr. Fengbiao Mao (University of Michigan, USA), for the advice on data visualization; Dr. Pierangelo Luporini (University of Camerino, Italy) and Dr. Estienne Swart (Max Planck Institute, Germany) for the advice on the preparation of the manuscript. This work was supported by the Aoshan Science and Technology Innovation Program of the Qingdao National Laboratory for Marine Science and Technology, National Natural Science Foundation of China (project No. 31772428), Young Elite Scientists Sponsorship Program by CAST (2017QNRC001) and the Fundamental Research Funds for the Central Universities (201841013 and 201762017). Research reported in this publication was also supported by the grants of the National Institutes of Health (award No. P40OD010964) and the National Science Foundation (grant No. 1158346) to N.A.S. and two grants to L.A.K. (NSF DEB-1541511 and NIH 1R15GM113177-01). The content is solely the responsibility of the authors and does not necessarily represent the official views of the National Institutes of Health. Any opinions, findings, and conclusions or recommendations expressed in this material are those of the author(s) and do not necessarily reflect the views of the National Science Foundation.

## AUTHOR CONTRIBUTION

X.C. and F.G. conceived the study; Y.J. and W.Z. provided the biological materials; X.C. designed the experiments; Y.J. performed the experiments; X.C. performed computational and experimental analysis for all figures and tables; X.C., F.G., C.L., L.A.K. and W.S. interpreted the data; N.A.S. and T.J.K. constructed the genome database website; X.C. wrote the paper with contribution from all authors. All authors read and approved the final manuscript.

## DATA AVAILABILITY

*Euplotes vannus* MAC genome assembly and gene annotation data including coding regions and predicted protein sequences are available at *Euplotes vannus* DB (EVDB, http://evan.ciliate.org).

## SUPPORTING INFORMATION

**Figure S1.** K-mer analysis of *Euplotes vannus* mating types to estimated genome size.

**Figure S2.** Genome assembly completeness evaluation of ciliates by BUSCO.

**Figure S3.** Venn diagram shows the genes annotated by BLASTX and Interproscan.

**Figure S4.** Schematic representation of the exon/intron boundaries with WebLogo in all 78661 introns in *E. vannus* mating type EVJ. The GTR and YAG motifs are well conserved.

**Figure S5.** A schema illustrates the criteria for detecting +1 frameshifting events. Blue boxes indicate the different BLASTX hits of a CDS region to a same target protein sequence (E-value cutoff = 1e-5). Grey boxes indicate the adjacent region between two BLASTX hits of a CDS region (inner distance cutoff = 10 nt). The brackets above denote the 0-frame codons and the brackets underneath denote the +1-frame codons. Yellow dots denote the nucleotides while the red ones denote the slippery site where frameshifting events occur.

**Figure S6.** Sequence alignment of the reverse complements of the MAC contigs containing Type-II pheromone coding genes in *E. vannus* and *E. crassus*. Blue and red boxes denote the start and stop codon of the coding region of pheromone genes, respectively.

**Figure S7.** Sequence alignment of the Type-I pheromone protein sequences in *E. vannus* and *E. crassus*. Identical residues are shadowed in black and similar residues are shaded in grey. Asterisks mark the positions of stop codons. Filled and light arrowheads indicate the extension positions of the pre- and pro-regions, respectively. Red dots denote the conserved cysteine residues in the secreted region. Numbers indicate the progressive amino acid positions in the sequences.

**Figure S8.** Sequence alignment of the reverse complements of the MAC contigs containing Type-I pheromone coding genes in *E. vannus* and *E. crassus*. Blue and red boxes denote the start and stop codon of the coding region of pheromone genes, respectively.

**Figure S9.** GBrowse snapshots of genomic and transcriptomic reads mapping on pheromone gene-related chromosomes in different mating types.

**Figure S10.** Overall gene expression level in starved or vegetative cells of different mating types.

**Figure S11.** Venn diagram shows a large part of mating type-specific transcripts is also starvation induced.

**Figure S12.** Species relationship and functional annotation of the mating type-specific and starvation induced transcripts.

**Figure S13.** Differential gene expression analysis under temperature, salinity and free ammonia stresses (relative to Figure 5). (A) PCA analysis on gene expression of mating type EVJ under different stresses. (B) Different environmental stresses activated or deactivated different gene groups.

**Figure S14.** Sequence alignment of Hsp70 protein sequences.

**Table S1.** Mating pattern observed when cultures of two mating types are mixed and genomic and transcriptomic (mRNA) data accessibility of six mating types of *E. vannus*.

**Table S2.** Genome assembly information of four mating types of *Euplotes vannus*.

**Table S3.** Genomic and transcriptomic reads mapping information.

**Table S4.** List of identified ncRNAs.

**Table S5.** Annotation information of repeats in the merged genome assembly of *E. vannus*.

**Table S6.** Expression and annotation information of *Euplotes vannus* genes.

**Table S7.** Homologs of mating-type loci in each mating type.

**Table S8.** Homologous search results by BLASTN reveal the relationship between coding regions of four pheromone genes in MAC genome and the corresponding MDS regions in MIC genome of *E. vannus*.

**Table S9.** PCR primers for genotyping of Type-II pheromone genes in *E. vannus* and determine mating types.

**Table S10.** Information of accession of genome/transcriptome assemblies of 32 ciliates.

## REFERENCES

Acorn AR, Clark KF, Jones S, et al. (2011) Analysis of expressed sequence tags (ESTs) and gene expression changes under different growth conditions for the ciliate *Anophryoides haemophila*, the causative agent of bumper car disease in the American lobster (*Homarus americanus*). Journal of Invertebrate Pathology, 107, 146–154. https://doi.org/10.1016/j.jip.2011.04.006

Aeschlimann SH, Jonsson F, Postberg J, et al. (2014) The draft assembly of the radically organized *Stylonychia lemnae* macronuclear genome. Genome Biol Evol, 6, 1707–1723. https://doi.org/10.1093/gbe/evu139

Akada R (1985) Mating types and mating-inducing factors (gamones) in the ciliate *Euplotes patella* syngen 2. Genetical Research, 46, 125–132. https://doi.org/10.1017/S0016672300022618

Alimenti C, Vallesi A, Federici S, et al. (2011) Isolation and structural characterization of two waterborne pheromones from *Euplotes crassus*, a ciliate commonly known to carry membrane-bound pheromones. Journal of Eukaryotic Microbiology, 58, 234–241. https://doi.org/10.1111/j.1550-7408.2011.00535.x

Bankevich A, Nurk S, Antipov D, et al. (2012) SPAdes: a new genome assembly algorithm and its applications to single-cell sequencing. Journal of Computational Biology, 19, 455–477. https://doi.org/10.1089/cmb.2012.0021

Borror AC (1980) Spatial distribution of marine ciliates: Micro-Ecologic and Biogeographic aspects of Protozoan Ecology. Journal of Eukaryotic Microbiology, 27, 10–13. https://doi.org/10.1111/j.1550-7408.1980.tb04224.x

Brown L, Mronga S, Bradshaw R, et al. (1993) Nuclear magnetic resonance solution structure of the pheromone Er-10 from the ciliated protozoan *Euplotes raikovi*. Journal of Molecular Biology, 231, 800–816. https://doi.org/10.1006/jmbi.1993.1327

Brünen-Nieweier C, Weiligmann JC, Hansen B, et al. (1998) The pheromones and pheromone genes of new stocks of the *Euplotes octocarinatus* species complex. European Journal of Protistology, 34, 124–132.

Burge SW, Daub J, Eberhardt R, et al. (2013) Rfam 11.0: 10 years of RNA families. Nucleic Acids Research, 41, 3. https://doi.org/10.1093/nar/gks1005

Camacho C, Coulouris G, Avagyan V, et al. (2009) BLAST+: architecture and applications. BMC Bioinformatics, 10, 1471–2105. https://doi.org/10.1186/1471-2105-10-421

Cervantes MD, Hamilton EP, Xiong J, et al. (2013) Selecting one of several mating types through gene segment joining and deletion in Tetrahymena thermophila. PLoS biology, 11, e1001518. https://doi.org/10.1371/journal.pbio.1001518

Chen X, Wang Y, Sheng Y, Warren A, Gao S (2018) GPSit: An automated method for evolutionary analysis of nonculturable ciliated microeukaryotes. Molecular Ecology Resources, 4, 1–14. https://doi.org/10.1111/1755-0998.12750

Chenna R, Sugawara H, Koike T, et al. (2003) Multiple sequence alignment with the Clustal series of programs. Nucleic Acids Research, 31, 3497–3500. https://doi.org/10.1093/nar/gkg500

Cheung P, Nigrelli R, Ruggieri G (1980) Studies on the morphology of *Uronema marinum* Dujardin (Ciliatea: Uronematidae) with a description of the histopathology of the infection in marine fishes. Journal of Fish Diseases, 3, 295–303. https://doi.org/10.1111/j.1365-2761.1980.tb00400.x

Clark MS, Peck LS (2009) HSP70 heat shock proteins and environmental stress in Antarctic marine organisms: a mini-review. Marine genomics, 2, 11–18. https://doi.org/10.1016/j.margen.2009.03.003

Crooks GE, Hon G, Chandonia JM, Brenner SE (2004) WebLogo: a sequence logo generator. Genome Research, 14, 1188–1190. https://doi.org/10.1101/gr.849004

Day JG, Gong Y, Hu Q (2017) Microzooplanktonic grazers - A potentially devastating threat to the commercial success of microalgal mass culture. Algal Research, 27, 356–365. https://doi.org/10.1016/j.algal.2017.08.024

Dhanker R, Kumar R, Tseng L-C, Hwang J-S (2013) Ciliate (*Euplotes* sp.) predation by *Pseudodiaptomus annandalei* (Copepoda: Calanoida) and the effects of mono-algal and pluri-algal diets. Zoological Studies, 52, 34. https://doi.org/10.1186/1810-522x-52-34

Di Giuseppe G, Erra F, Dini F, et al. (2011) Antarctic and Arctic populations of the ciliate *Euplotes nobilii* show common pheromone-mediated cell-cell signaling and cross-mating. Proceedings of the National Academy of Sciences of the United States of America, 108, 3181–3186. https://doi.org/10.1073/pnas.1019432108

Dini F, Luporini P (1985) Mating-type polymorphic variation in *Euplotes minuta* (Ciliophora: Hypotrichida). The Journal of protozoology, 32, 111–117.

Dini F, Nyberg D (1993) Sex in ciliates. In: Advances in Microbial Ecology (ed. Nelson KE), pp. 85–153. Springer.

Dobzhansky T (1940) Speciation as a stage in evolutionary divergence. The American Naturalist, 74, 312–321. https://doi.org/10.1086/280899

Dragesco A, Dragesco J, Coste F, et al. (1995) *Philasterides dicentrarchi*, n. sp.,(Ciliophora, Scuticociliatida), a histophagous opportunistic parasite of *Dicentrarchus labrax* (Linnaeus, 1758), a reared marine fish. European Journal of Protistology, 31, 327–340. https://doi.org/10.1016/S0932-4739(11)80097-0

Edgar RC (2004) MUSCLE: multiple sequence alignment with high accuracy and high throughput. Nucleic Acids Research, 32, 1792–1797. https://doi.org/10.1093/nar/gkh340

Fernandes M (1994) Structure and regulation of heat shock gene promoters. In: The Biology of Heat Shock Proteins and Molecular Chaperones, pp. 375–393. Cold Spring Harbor Laboratory Press.

Foissner W (2006) Biogeography and dispersal of micro-organisms: a review emphasizing protists. Acta Protozoologica, 45, 111–136.

Foissner W, Chao A, Katz LA (2008) Diversity and geographic distribution of ciliates (Protista: Ciliophora). Biodiversity and Conservation, 17, 345–363. https://doi.org/10.1007/s10531-007-9254-7

Frazee AC, Pertea G, Jaffe AE, et al. (2015) Ballgown bridges the gap between transcriptome assembly and expression analysis. Nature Biotechnology, 33, 243–246. https://doi.org/10.1038/nbt.3172.

Fu L, Niu B, Zhu Z, Wu S, Li W (2012) CD-HIT: accelerated for clustering the next-generation sequencing data. Bioinformatics, 28, 3150–3152. https://doi.org/10.1093/bioinformatics/bts565

Gentekaki E, Kolisko M, Gong Y, Lynn D (2017) Phylogenomics solves a long-standing evolutionary puzzle in the ciliate world: The subclass Peritrichia is monophyletic. Molecular Phylogenetics and Evolution, 106, 1–5. https://doi.org/10.1016/j.ympev.2016.09.016

Gong J, Song W, Warren A (2005) Periphytic ciliate colonization: annual cycle and responses to environmental conditions. Aquatic Microbial Ecology, 39, 159–170. https://doi.org/10.3354/ame039159

Gutiérrez JC, Martín-González A, Díaz S, Ortega R (2003) Ciliates as a potential source of cellular and molecular biomarkers/biosensors for heavy metal pollution. European Journal of Protistology, 39, 461–467. https://doi.org/10.1078/0932-4739-00021

Hadjivasiliou Z, Iwasa Y, Pomiankowski A (2015) Cell-cell signalling in sexual chemotaxis: a basis for gametic differentiation, mating types and sexes. J R Soc Interface, 12, 20150342. https://doi.org/10.1098/rsif.2015.0342

Heaphy SM, Mariotti M, Gladyshev VN, Atkins JF, Baranov PV (2016) Novel ciliate genetic code variants including the reassignment of all three stop codons to sense codons in *Condylostoma magnum*. Molecular Biology and Evolution, 33, 2885–2889. https://doi.org/10.1093/molbev/msw166

Heckmann K (1964) Experimentelle Untersuchungen an *Euplotes crassus*. Zeitschrift für Vererbungslehre, 95, 114–124. https://doi.org/10.1007/Bf00894912

Heckmann K, Kuhlmann HW (1986) Mating types and mating inducing substances in *Euplotes octocarinatus*. Journal of Experimental Zoology, 237, 87–96. https://doi.org/10.1002/jez.1402370113

Huang X, Madan A (1999) CAP3: A DNA sequence assembly program. Genome Research, 9, 868–877.

Iglesias R, Paramá A, Alvarez M, et al. (2001) *Philasterides dicentrarchi* (Ciliophora, Scuticociliatida) as the causative agent of scuticociliatosis in farmed turbot *Scophthalmus maximus* in Galicia (NW Spain). Diseases of Aquatic Organisms, 46, 47–55. https://doi.org/10.3354/dao046047

Jerka-Dziadosz M, Dosche C, Kuhlmann HW, Heckmann K (1987) Signal-induced reorganization of the microtubular cytoskeleton in the ciliated protozoon *Euplotes octocarinatus*. Journal of Cell Science, 87, 555–564.

Jiang L, Morin PJ (2004) Temperature-dependent interactions explain unexpected responses to environmental warming in communities of competitors. Journal of Animal Ecology, 73, 569–576. https://doi.org/10.1111/j.0021-8790.2004.00830.x

Jiang Y, Xu H, Hu X, et al. (2011) An approach to analyzing spatial patterns of planktonic ciliate communities for monitoring water quality in Jiaozhou Bay, northern China. Marine Pollution Bulletin, 62, 227–235. https://doi.org/10.1016/j.marpolbul.2010.11.008

Jones P, Binns D, Chang HY, et al. (2014) InterProScan 5: genome-scale protein function classification. Bioinformatics, 30, 1236–1240. https://doi.org/10.1093/bioinformatics/btu031

Keeling PJ, Burki F, Wilcox HM, et al. (2014) The Marine Microbial Eukaryote Transcriptome Sequencing Project (MMETSP): illuminating the functional diversity of eukaryotic life in the oceans through transcriptome sequencing. PLoS Biol, 12. https://doi.org/10.1371/journal.pbio.1001889

Kim B-M, Rhee J-S, Choi I-Y, Lee Y-M (2018) Transcriptional profiling of antioxidant defense system and heat shock protein (Hsp) families in the cadmium-and copper-exposed marine ciliate *Euplotes crassus*. Genes & Genomics, 40, 85–98. https://doi.org/10.1007/s13258-017-0611-y

Kim D, Langmead B, Salzberg SL (2015) HISAT: a fast spliced aligner with low memory requirements. Nature Methods, 12, 357–360. https://doi.org/10.1038/nmeth.3317

Kim S-J, Kim J-H, Ju S-J (2017) Adaptation responses of individuals to environmental changes in the ciliate *Euplotes crassus*. Ocean Science Journal, 52, 127–138. https://doi.org/10.1007/s12601-017-0014-7

Kimball R (1942) The nature and inheritance of mating types in *Euplotes patella*. Genetics, 27, 269.

Kobayashi N, McENTEE K (1993) Identification of cis and trans components of a novel heat shock stress regulatory pathway in *Saccharomyces cerevisiae*. Molecular and Cellular Biology, 13, 248–256.

Kohl M, Wiese S, Warscheid B (2011) Cytoscape: software for visualization and analysis of biological networks. Methods in Molecular Biology, 696, 291–303. https://doi.org/10.1007/978-1-60761-987-1_18

Kuhlmann H-W (1994) Escape response of *Euplotes octocarinatus* to turbellarian predators. Archiv für Protistenkunde, 144, 163–171. https://doi.org/10.1016/S0003-9365(11)80121-1

Kuhlmann H-W, Brünen-Nieweler C, Heckmann K (1997) Pheromones of the ciliate *Euplotes octocarinatus* not only induce conjugation but also function as chemoattractants. Journal of Experimental Zoology, 277, 38–48.

Kuhlmann H-W, Heckmann K (1994) Predation risk of typical ovoid and ’winged’ morphs of *Euplotes* (Protozoa, Ciliophora). Hydrobiologia, 284, 219–227. https://doi.org/10.1007/Bf00006691

Kumar S, Stecher G, Tamura K (2016) MEGA7: Molecular Evolutionary Genetics Analysis version 7.0 for bigger datasets. Molecular Biology and Evolution, 33, 1870–1874. https://doi.org/10.1093/molbev/msw054

Kusch J (1993a) Behavioural and morphological changes in ciliates induced by the predator *Amoeba proteus*. Oecologia, 96, 354–359. https://doi.org/10.1007/BF00317505

Kusch J (1993b) Induction of defensive morphological changes in ciliates. Oecologia, 94, 571–575. https://doi.org/10.1007/BF00566974

Kusch J (1995) Adaptation of inducible defense in *Euplotes daidaleos* (Ciliophora) to predation risks by various predators. Microbial Ecology, 30, 79–88.

Kusch J, Kuhlmann HW (1994) Cost of Stenostomum-induced morphological defence in the ciliate *Euplotes octocarinatus*. Archiv Fur Hydrobiologie, 130, 257–267.

La Terza A, Miceli C, Luporini P (2004) The gene for the heat-shock protein 70 of *Euplotes focardii*, an Antarctic psychrophilic ciliate. Antarctic Science, 16, 23–28. https://doi.org/10.1017/S0954102004001774

La Terza A, Papa G, Miceli C, Luporini P (2001) Divergence between two Antarctic species of the ciliate *Euplotes, E. focardii* and *E. nobilii*, in the expression of heat-shock protein 70 genes. Molecular Ecology, 10, 1061–1067. https://doi.org/10.1046/j.1365-294X.2001.01242.x

La Terza A, Passini V, Barchetta S, Luporini P (2007) Adaptive evolution of the heat-shock response in the Antarctic psychrophilic ciliate, *Euplotes focardii*: hints from a comparative determination of the hsp70 gene structure. Antarctic Science, 19, 239–244. https://doi.org/10.1017/S0954102007000314

Langfelder P, Horvath S (2008) WGCNA: an R package for weighted correlation network analysis. BMC Bioinformatics, 9, 559. https://doi.org/10.1186/1471-2105-9-559

Le SQ, Gascuel O (2008) An improved general amino acid replacement matrix. Molecular Biology and Evolution, 25, 1307–1320. https://doi.org/10.1093/molbev/msn067

Liao Y, Smyth GK, Shi W (2013) featureCounts: an efficient general purpose program for assigning sequence reads to genomic features. Bioinformatics, 30, 923–930. https://doi.org/10.1093/bioinformatics/btt656

Liu A, Luginbühl P, Zerbe O, et al. (2001) NMR structure of the pheromone Er-22 from *Euplotes raikovi*. Journal of Biomolecular NMR, 19, 75–78.

Liu W, Jiang J, Xu Y, et al. (2017) Diversity of free-living marine ciliates (Alveolata, Ciliophora): faunal studies in coastal waters of China during the years 2011-2016. European Journal of Protistology, 61, 424–438. https://doi.org/10.1016/j.ejop.2017.04.007

Lobanov AV, Heaphy SM, Turanov AA, et al. (2017) Position-dependent termination and widespread obligatory frameshifting in *Euplotes* translation. Nature Structural & Molecular Biology, 24, 61–68. https://doi.org/10.1038/nsmb.3330

Love MI, Huber W, Anders S (2014) Moderated estimation of fold change and dispersion for RNA-seq data with DESeq2. Genome Biology, 15, 014–0550. https://doi.org/10.1186/s13059-014-0550-8

Lowe TM, Eddy SR (1997) tRNAscan-SE: a program for improved detection of transfer RNA genes in genomic sequence. Nucleic Acids Research, 25, 955–964.

Luginbühl P, Ottiger M, Mronga S, WÜthrich K (1994) Structure comparison of the pheromones Er-1, Er-10, and Er-2 from *Euplotes raikovi*. Protein Science, 3, 1537–1546. https://doi.org/10.1002/pro.5560030919

Luo X, Gao F, Yi Z, et al. (2017) Taxonomy and molecular phylogeny of two new brackish hypotrichous ciliates, with the establishment of a new genus (Ciliophora, Spirotrichea). Zoological Journal of The Linnean Society, 179, 475–491. https://doi.org/10.1111/zoj.12451

Lynn D (2009) Ciliates. In: Encyclopedia of Microbiology (Third Edition) (ed. Schaechter M), pp. 578–592. Academic Press, Oxford.

Maere S, Heymans K, Kuiper M (2005) BiNGO: a Cytoscape plugin to assess overrepresentation of gene ontology categories in biological networks. Bioinformatics, 21, 3448–3449. https://doi.org/10.1093/bioinformatics/bti551

Möllenbeck M, Heckmann K (1999) Characterization of two genes encoding a fifth so far unknown pheromone of *Euplotes octocarinatus*. European Journal of Protistology, 35, 225–230.

Morshauser RC, Hu W, Wang H, et al. (1999) High-resolution solution structure of the 18 kDa substrate-binding domain of the mammalian chaperone protein Hsc701. Journal of Molecular Biology, 289, 1387–1403. https://doi.org/10.1006/jmbi.1999.2776

Mronga S, Luginbühl P, Brown L, et al. (1994) The NMR solution structure of the pheromone Er-1 from the ciliated protozoan *Euplotes raikovi*. Protein Science, 3, 1527–1536. https://doi.org/10.1002/pro.5560030918

Munday B, O’Donoghue P, Watts M, Rough K, Hawkesford T (1997) Fatal encephalitis due to the scuticociliate *Uronema nigricans* in sea-caged, southern bluefin tuna *Thunnus maccoyii*. Diseases of Aquatic Organisms, 30, 17–25. https://doi.org/10.3354/dao030017

Nanney D, Caughey PA, Tefankjian A (1955) The genetic control of mating type potentialities in *Tetrahymena pyriformis*. Genetics, 40, 668.

Nobili R, Luporini P, Dini F (1978) Breeding systems, species relationships and evolutionary trends in some marine species of Euplotidae (Hypotrichida Ciliata). Marine organisms: genetics, ecology, and evolution. Plenum, New York, 591–616.

Nurk S, Bankevich A, Antipov D, et al. (2013) Assembling single-cell genomes and mini-metagenomes from chimeric MDA products. Journal of Computational Biology, 20, 714–737. https://doi.org/10.1089/cmb.2013.0084

Orias E, Singh DP, Meyer E (2017) Genetics and Epigenetics of mating type determination in *Paramecium* and *Tetrahymena*. Annual Review of Microbiology, 71, 133–156. https://doi.org/10.1146/annurev-micro-090816-093342

Ottiger M, Szyperski T, Luginbühl P, et al. (1994) The NMR solution structure of the pheromone Er-2 from the ciliated protozoan *Euplotes raikovi*. Protein Science, 3, 1515–1526. https://doi.org/10.1002/pro.5560030917

Pedrini B, Placzek WJ, Koculi E, et al. (2007) Cold-adaptation in sea-water-borne signal proteins: sequence and NMR structure of the pheromone En-6 from the Antarctic ciliate *Euplotes nobilii*. Journal of Molecular Biology, 372, 277–286. https://doi.org/10.1016/j.jmb.2007.06.046

Persoone G, Uyttersprot G (1975) The influence of inorganic and organic pollutants on the rate of reproduction of a marine hypotrichous ciliate: *Euplotes vannus* Muller. Revue Internationale d’Oceanographie Medicale, 37, 125–152.

Pertea M, Pertea GM, Antonescu CM, et al. (2015) StringTie enables improved reconstruction of a transcriptome from RNA-seq reads. Nature Biotechnology, 33, 290–295. https://doi.org/10.1038/nbt.3122

Petz W, Valbonesi A, Schiftner U, Quesada A, Cynan Ellis-Evans J (2007) Ciliate biogeography in Antarctic and Arctic freshwater ecosystems: endemism or global distribution of species? Fems Microbiology Ecology, 59, 396–408. https://doi.org/10.1111/j.1574-6941.2006.00259.x

Placzek WJ, Etezady-Esfarjani T, Herrmann T, et al. (2007) Cold-adapted signal proteins: NMR structures of pheromones from the antarctic ciliate *Euplotes nobilii*. Iubmb Life, 59, 578–585. https://doi.org/10.1080/15216540701258165

Ruis H, Schüller C (1995) Stress signaling in yeast. Bioessays, 17, 959–965. https://doi.org/10.1002/bies.950171109

Schulze Dieckhoff H, Freiburg M, Heckmann K (1987) The isolation of gamones 3 and 4 of *Euplotes octocarinatus*. The FEBS Journal, 168, 89–94.

Siegel R, Larison L (1960) The genic control of mating types in *Paramecium bursaria*. Proceedings of the National Academy of Sciences of the United States of America, 46, 344–349.

Simao FA, Waterhouse RM, Ioannidis P, Kriventseva EV, Zdobnov EM (2015) BUSCO: assessing genome assembly and annotation completeness with single-copy orthologs. Bioinformatics, 31, 3210–3212. https://doi.org/10.1093/bioinformatics/btv351

Singh DP, Saudemont B, Guglielmi G, et al. (2014) Genome-defence small RNAs exapted for epigenetic mating-type inheritance. Nature, 509, 447. https://doi.org/10.1038/nature13318

Slabodnick MM, Ruby JG, Reiff SB, et al. (2017) The macronuclear genome of *Stentor coeruleus* reveals tiny introns in a giant cell. Current Biology, 27, 569–575. https://doi.org/10.1016/j.cub.2016.12.057

Sriram M, Osipiuk J, Freeman B, Morimoto R, Joachimiak A (1997) Human Hsp70 molecular chaperone binds two calcium ions within the ATPase domain. Structure, 5, 403–414. https://doi.org/10.1016/S0969-2126(97)00197-4

Stamatakis A (2014) RAxML version 8: a tool for phylogenetic analysis and post-analysis of large phylogenies. Bioinformatics, 30, 1312–1313. https://doi.org/10.1093/bioinformatics/btu033

Stanke M, Tzvetkova A, Morgenstern B (2006) AUGUSTUS at EGASP: using EST, protein and genomic alignments for improved gene prediction in the human genome. Genome Biology, 7, 1. https://doi.org/10.1186/gb-2006-7-s1-s11

Stein LD (2013) Using GBrowse 2.0 to visualize and share next-generation sequence data. Briefings in Bioinformatics, 14, 162–171. https://doi.org/10.1093/bib/bbt001

Stoeck T, Kochems R, Forster D, Lejzerowicz F, Pawlowski J (2018) Metabarcoding of benthic ciliate communities shows high potential for environmental monitoring in salmon aquaculture. Ecological Indicators, 85, 153–164. https://doi.org/10.1016/j.ecolind.2017.10.041

Swart EC, Bracht JR, Magrini V, et al. (2013) The *Oxytricha trifallax* macronuclear genome: a complex eukaryotic genome with 16,000 tiny chromosomes. PLoS Biol, 11, 29. https://doi.org/10.1371/journal.pbio.1001473

Swart EC, Serra V, Petroni G, Nowacki M (2016) Genetic codes with no dedicated stop codon: context-dependent translation termination. Cell, 166, 691–702. https://doi.org/10.1016/j.cell.2016.06.020

Tarailo-Graovac M, Chen N (2009) Using RepeatMasker to identify repetitive elements in genomic sequences. Current Protocols in Bioinformatics, 4, 4–10. https://doi.org/10.1002/0471250953.bi0410s25

Trielli F, Amaroli A, Sifredi F, et al. (2007) Effects of xenobiotic compounds on the cell activities of *Euplotes crassus*, a single-cell eukaryotic test organism for the study of the pollution of marine sediments. Aquatic Toxicology, 83, 272–283. https://doi.org/10.1016/j.aquatox.2007.05.002

Valbonesi A, Ortenzi C, Luporini P (1988) An integrated study of the species problem in the *Euplotes crassus-minuta-vannus* group. Journal of Eukaryotic Microbiology, 35, 38–45. https://doi.org/10.1111/j.1550-7408.1988.tb04073.x

Vallesi A, Alimenti C, Federici S, et al. (2014) Evidence for gene duplication and allelic codominance (not hierarchical dominance) at the mating-type locus of the ciliate, *Euplotes crassus*. Journal of Eukaryotic Microbiology, 61, 620–629. https://doi.org/10.1111/jeu.12140

Vallesi A, Alimenti C, Luporini P (2016) Ciliate pheromones: primordial self-/nonself-recognition signals. In: Lessons in Immunity (eds. Ballarin L, Cammarata M), pp. 1–16. Elsevier.

Vallesi A, Alimenti C, Pedrini B, et al. (2012) Coding genes and molecular structures of the diffusible signalling proteins (pheromones) of the polar ciliate, *Euplotes nobilii*. Marine genomics, 8, 9–13. https://doi.org/10.1016/j.margen.2012.03.004

Walton BM, Gates MA, Kloos A, Fisher J (1995) Intraspecific variability in the thermal dependence of locomotion, population growth, and mating in the ciliated protist *Euplotes vannus*. Physiological Zoology, 68, 98–113. https://doi.org/10.1086/physzool.68.1.30163920

Wang C, Zhang T, Wang Y, et al. (2017a) Disentangling sources of variation in SSU rDNA sequences from single cell analyses of ciliates: impact of copy number variation and experimental error. Proceedings of the Royal Society B: Biological Sciences, 284, 20170425. https://doi.org/10.1098/rspb.2017.0425

Wang R, Miao W, Wang W, Xiong J, Liang A (2018) EOGD: the *Euplotes octocarinatus* genome database. BMC Genomics, 19, 63. https://doi.org/10.1186/s12864-018-4445-z

Wang R, Xiong J, Wang W, Miao W, Liang A (2016) High frequency of +1 programmed ribosomal frameshifting in *Euplotes octocarinatus*. Scientific Reports, 6, 21139. https://doi.org/10.1038/srep21139

Wang Y, Chen X, Sheng Y, Liu Y, Gao S (2017b) N6-adenine DNA methylation is associated with the linker DNA of H2A. Z-containing well-positioned nucleosomes in Pol II-transcribed genes in *Tetrahymena*. Nucleic Acids Research, 45, 11594–11606. https://doi.org/10.1093/nar/gkx883

Wang Y, Wang Y, Sheng Y, et al. (2017c) A comparative study of genome organization and epigenetic mechanisms in model ciliates, with an emphasis on *Tetrahymena, Paramecium* and *Oxytricha*. European Journal of Protistology, 61, 376–387. https://doi.org/10.1016/j.ejop.2017.06.006

Weiss MS, Anderson DH, Raffioni S, et al. (1995) A cooperative model for receptor recognition and cell adhesion: evidence from the molecular packing in the 1.6-A crystal structure of the pheromone Er-1 from the ciliated protozoan *Euplotes raikovi*. Proceedings of the National Academy of Sciences of the United States of America, 92, 10172–10176. https://doi.org/10.1073/pnas.92.22.10172

Wiąckowski K, Szkarłat M (1996) Effects of food availability on predator-induced morphological defence in the ciliate *Euplotes octocarinatus* (Protista). Hydrobiologia, 321, 47–52. https://doi.org/10.1007/Bf00018676

Wickham H (2016) ggplot2: elegant graphics for data analysis Springer.

Xiong J, Gao S, Dui W, et al. (2016) Dissecting relative contributions of cis-and trans-determinants to nucleosome distribution by comparing *Tetrahymena* macronuclear and micronuclear chromatin. Nucleic Acids Research, 44, 10091–10105. https://doi.org/10.1093/nar/gkw684

Xu H, Song W, Warren A (2004) An investigation of the tolerance to ammonia of the marine ciliate *Euplotes vannus* (Protozoa, Ciliophora). Hydrobiologia, 519, 189–195. https://doi.org/10.1023/B:HYDR.0000026505.91684.ab

Xu H, Zhang W, Jiang Y, Yang EJ (2014) Use of biofilm-dwelling ciliate communities to determine environmental quality status of coastal waters. Science of The Total Environment, 470, 511–518. https://doi.org/10.1016/j.scitotenv.2013.10.025

Yan Y, Rogers AJ, Gao F, Katz LA (2017) Unusual features of non-dividing somatic macronuclei in the ciliate class Karyorelictea. European Journal of Protistology, 61, 399–408. https://doi.org/10.1016/j.ejop.2017.05.002

Yeomans W, Chubb J, Sweeting R (1997) Use of protozoan communities for pollution monitoring. Parassitologia, 39, 201–212.

Zahn R, Damberger F, Ortenzi C, Luporini P, Wüthrich K (2001) NMR structure of the *Euplotes raikovi* pheromone Er-23 and identification of its five disulfide bonds. Journal of Molecular Biology, 313, 923–931. https://doi.org/10.1006/jmbi.2001.5099

Zhao Y, Yi Z, Warren A, Song WB (2018) Species delimitation for the molecular taxonomy and ecology of the widely distributed microbial eukaryote genus *Euplotes* (Alveolata, Ciliophora). Proceedings of the Royal Society B: Biological Sciences, 285, 20172159. https://doi.org/10.1098/rspb.2017.2159

